# *Drosophila* Histone Locus Body assembly and function involves multiple interactions

**DOI:** 10.1101/2020.03.16.994483

**Authors:** Kaitlin P. Koreski, Leila E. Rieder, Lyndsey M. McLain, William F. Marzluff, Robert J. Duronio

## Abstract

The histone locus body (HLB) assembles at replication-dependent (RD) histone loci and concentrates factors required for RD histone mRNA biosynthesis. The *D. melanogaster* genome has a single locus comprised of ∼100 copies of a tandemly arrayed repeat unit containing one copy of each of the 5 RD histone genes. To determine sequence elements required for *D. melanogaster* HLB formation and histone gene expression, we used transgenic gene arrays containing 12 copies of the histone repeat unit that functionally complement loss of the ∼200 endogenous RD histone genes. A 12x histone gene array in which all *H3-H4* promoters were replaced with *H2a-H2b* promoters does not form an HLB or express high levels of RD histone mRNA in the presence of the endogenous histone genes. In contrast, this same transgenic array is active in HLB assembly and RD histone gene expression in the absence of the endogenous RD histone genes and rescues the lethality caused by homozygous deletion of the RD histone locus. The HLB formed in the absence of endogenous RD histone genes on the mutant 12x array contains all known factors present in the wild type HLB including CLAMP, which normally binds to GAGA repeats in the *H3-H4* promoter. These data suggest that multiple protein-protein and/or protein-DNA interactions contribute to HLB formation, and that the large number of endogenous RD histone gene copies sequester available factor(s) from attenuated transgenic arrays, thereby preventing HLB formation and gene expression.

## Introduction

An important organizing principle in cells is the use of membraneless compartments to spatially and temporally regulate diverse biological processes (Mitrea and Kriwacki, 2016). Numerous membraneless compartments have been identified in both the nucleus (e.g. nucleoli, Cajal bodies, histone locus bodies) and the cytoplasm (e.g. P-bodies, stress granules, germ granules) and are collectively referred to as biomolecular condensates (Banani *et al*., 2017). There is increasing evidence suggesting that membraneless compartments are formed through liquid-liquid phase separation or condensation (Alberti *et al*., 2019). This occurs when proteins and/or nucleic acids in the nucleoplasm or cytoplasm coalesce or demix into a condensed phase that often resembles liquid droplets. Large nuclear condensates that are visible under light microscopy are most often referred to as nuclear bodies (NBs), and represent an important organizing feature of the nucleus.

The histone locus body (HLB) is a conserved NB that assembles at replication-dependent (RD) histone genes and concentrates factors required for RD histone mRNA biogenesis (Duronio and Marzluff, 2017). RD histone mRNAs are the only eukaryotic mRNAs that are not polyadenylated (Marzluff and Koreski, 2017). The unique stem loop at the 3’end of RD histone mRNAs results from a processing reaction requiring a specialized suite of factors, some of which are constitutively localized in the HLB (Duronio and Marzluff, 2017). The HLB provides a powerful system to study how NBs form and function because it contains a well characterized set of factors involved in producing a unique class of cell-cycle regulated mRNAs. We previously demonstrated that concentrating factors (e.g. FLASH and U7 snRNP) in the *Drosophila melanogaster* HLB is critical for efficient histone pre-mRNA processing (Wagner *et al*., 2007; Tatomer *et al*., 2016b). However, a full understanding of how the HLB participates in histone mRNA biosynthesis requires knowledge of HLB assembly at the molecular level.

Prior studies of NBs have provided several important assembly concepts that are applicable to the HLB. Many NB components have an intrinsic ability to self-associate, an observation leading to two different models of NB assembly: (1) interactions among NB components occur stochastically, wherein individual factors can be recruited to the body in any order, or (2) components assemble in an ordered or hierarchical pathway, wherein the recruitment of components is predicated on prior recruitment of others (Dundr and Misteli, 2010; Rajendra *et al*., 2010). The HLB appears to employ a hybrid version of these two possibilities. For example, genetic loss of function experiments suggest a partially ordered assembly pathway of the *Drosophila* HLB with some components being required for subsequent recruitment of others (White *et al*., 2011; Terzo *et al*., 2015; Tatomer *et al*., 2016a). The scaffolding protein Mxc (Multi sex combs), the *Drosophila* ortholog of human NPAT (Nuclear Protein, Ataxia-Telangiectasia Locus), and FLASH (FLICE-Associated Huge protein) likely form the core of the HLB and are required for subsequent recruitment of other factors (White *et al*., 2011). Tethering experiments indicate that ectopic HLB formation also may be induced by several different HLB components, supporting a stochastic model of assembly (Shevtsov and Dundr, 2011).

The initiation event in self-organizing NB assembly is the key step in the process and is not well understood. A prevalent model postulates a “seeding” event that initiates the nucleation of critical components that form a platform for further recruitment of other components (Dundr, 2011; Shevtsov and Dundr, 2011; Altmeyer *et al*., 2015; Falahati *et al*., 2016; Stanek and Fox, 2017; Gomes and Shorter, 2019). In some instances, RNA is thought to help seed NB assembly, and NBs such as the nucleolus and ectopic paraspeckles can form at sites of specific transcription (Matera *et al*., 2009; Mao *et al*., 2011; Falahati *et al*., 2016). Blocking transcription prevents complete HLB assembly in both zebrafish and flies (Salzler *et al*., 2013; Heyn *et al*., 2017; Hur, 2019). However, the HLB is present at RD histone genes even in G1 when the genes are not active, raising the possibility that histone genes themselves participate in seeding HLB assembly (Zhao *et al*., 1998; Liu *et al*., 2006).

In *Drosophila*, the RD histone genes are present at a single locus with ∼100 copies of a tandemly arrayed 5kb repeat unit, each of which contains one copy of the divergently transcribed *H2a-H2b* and *H3-H4* gene pairs as well as the gene for linker histone *H1* (Lifton *et al*., 1978; McKay *et al*., 2015; Bongartz and Schloissnig, 2019). Using transgenes containing a wild type or mutant derivative of a single histone repeat, we previously demonstrated that the bidirectional *H3-H4* promoter stimulated HLB assembly and transcription of the single histone repeat in salivary glands (Salzler *et al*., 2013). We subsequently showed that the conserved GAGA repeat elements present in the *H3-H4* promoter region are targeted by the zinc-finger transcription factor CLAMP (Chromatin-Linked Adaptor for MSL Proteins), and that this interaction promotes HLB assembly (Rieder *et al*., 2017). Thus, the *H3-H4* promoter region might act to seed HLB assembly.

In this work, we leveraged transgenic histone gene arrays to test whether the *H3-H4* promoter region is necessary for *in vivo* function of the RD histone locus. We found that replacement of *H3-H4* promoters with *H2a-H2b* promoters results in an attenuated transgenic histone gene array that does not function in the presence of the intact endogenous RD histone locus, but surprisingly provides full *in vivo* function, including normal HLB assembly and histone gene expression, when the endogenous RD histone locus is absent. These results suggest that multiple elements in the histone genes and core HLB proteins are involved in HLB assembly.

## Results

To study histone gene regulation and the role of histone post-translational modifications in chromatin we previously developed an experimental system in which BAC-based, transgenic histone gene arrays containing 12 copies of the 5kb histone repeat unit assemble HLBs and functionally complement homozygous loss of the ∼100 gene copies at the endogenous RD histone locus, *HisC* (McKay *et al*., 2015). To study histone gene expression, we created a 12x^HWT^ (<u>H</u>istone <u>W</u>ild <u>T</u>ype) transgenic construct containing a synonymous polymorphism in the *H2a* gene (i.e. mutation of a XhoI site) that allows us to distinguish transgenic *H2a* gene expression from endogenous *H2a* gene expression (McKay *et al*., 2015). Here, we extended this design and created a “<u>D</u>esigner <u>W</u>ild <u>T</u>ype” (DWT) 5kb histone repeat unit that has all five histone genes marked in a similar manner (Fig. 1A, Fig. S1A). We also introduced restriction enzyme sites around the mRNA 3’ end processing signals, the stem loop and histone downstream element (Marzluff and Koreski, 2017), to allow us to readily manipulate those sequences (Fig. S1). We used this DWT repeat to create a 12x^DWT^ construct, and found that a transgenic version of this array behaved identically to the original 12x^HWT^ array. It rescued the lethality caused by homozygous deletion of *HisC*, resulting in viable, fertile adult flies. In addition, we can maintain a stock of flies lacking the endogenous RD histone locus and containing a homozygous 12x^DWT^ transgene.

**Figure 1.**
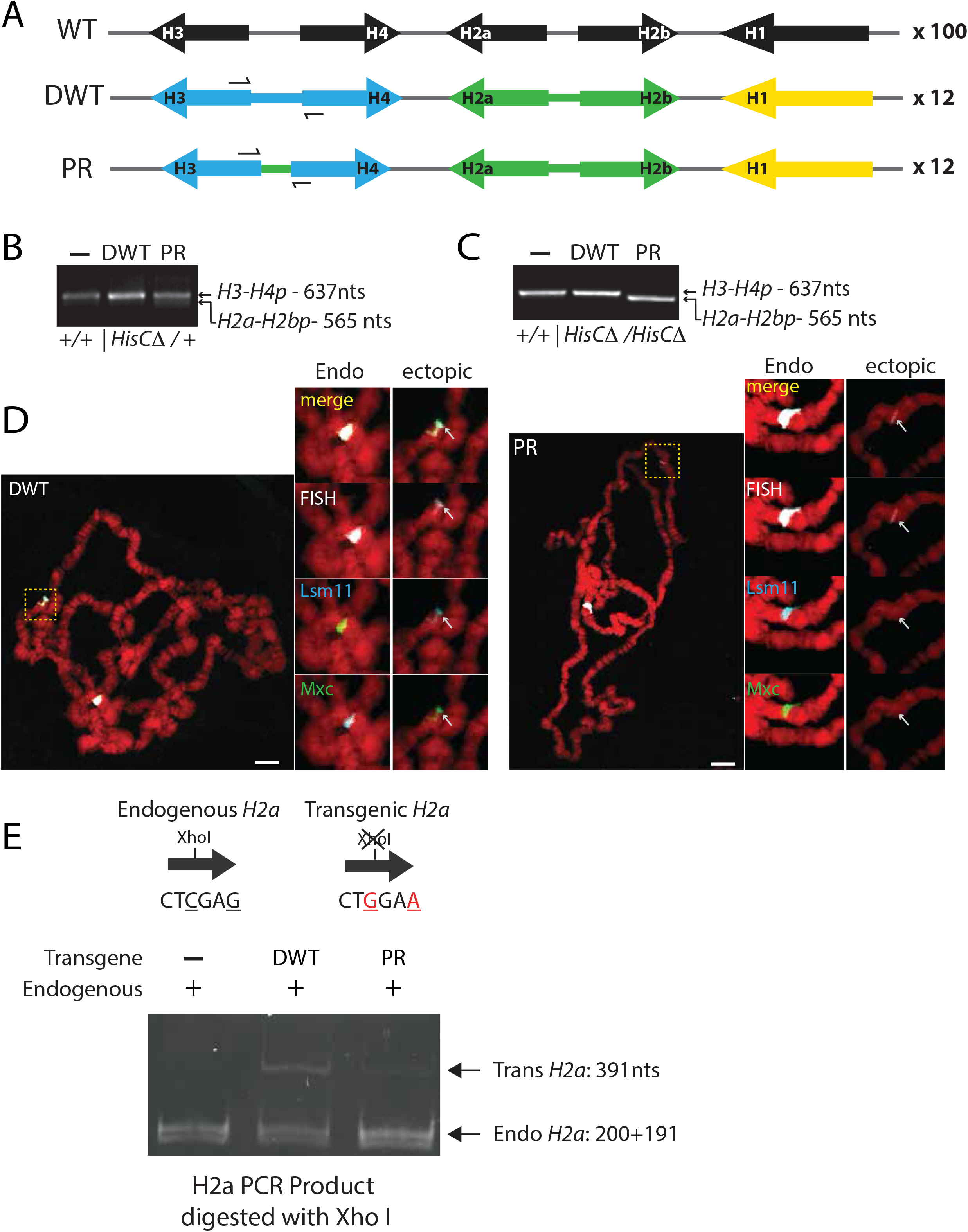
The *H3-H4* bidirectional promoter is required for ectopic HLB formation in the presence of the endogenous RD histone genes. **A)** Schematic of the endogenous *Drosophila* RD histone gene repeat, which is found in a tandem array of ∼100 copies at *HisC*, and the BAC-based DWT and PR synthetic histone repeats, which arrayed to 12 copies and inserted at VK33 on chromosome 3. The colors indicate that each transgenic histone gene was engineered to be distinguishable molecularly from the endogenous histone genes (see Fig. S1). In the PR construct the *H2a-H2b* promoters (green) replaced the *H3-H4* promoters (blue). **B, C)** Primers that anneal to *H3* and *H4* genes (A) were used to amplify the promoter region from genomic DNA extracted from whole flies heterozygous (B) or embryos homozygous (C) for a deletion of the *HisC* locus and containing either the 12x^DWT^ or 12x^PR^ transgene. The amplicon from a wild type locus is larger (637nts) than the PR locus (565nts). **D)** Polytene chromosome squashes from 3^rd^ instar larval salivary glands containing both *HisC* and either the 12x^DWT^ or 12x^PR^ transgenes hybridized with a probe to the histone locus and stained for HLB components Lsm11 and Mxc. Note that absence of an HLB at 12x^PR^ (arrows) when *HisC* is present (right panels). Scale bar 10 microns. **E)** Assay for specific detection of endogenous versus transgenic RD histone gene expression. Total RNA was prepared from 3^rd^ instar larvae containing the endogenous genes and one copy of the 12x^DWT^ or 12x^PR^ transgene. RT-PCR with *H2a* primers followed by digestion with *Xho I* cleaves cDNA from the endogenous genes but not the transgenes, which was visualized using an 8% polyacrylamide gel.

### The H3-H4 promoter stimulates HLB formation

To test whether the *H3-H4* promoter region is necessary for histone locus function in vivo, we engineered a derivative of the 12x^DWT^ array in which all *H3-H4* promoters were replaced with *H2a-H2b* promoters. In this “<u>P</u>romoter <u>R</u>eplacement” (12x^PR^) construct, we replaced the entire 298nt sequence between the initiation codons of the divergently transcribed *H3* and *H4* genes with the 226nt sequence between the initiation codons of the divergently transcribed *H2a* and *H2b* genes, leaving the *H1* and *H2a* and *H2b* genes intact (Fig. 1A, Fig. S1B). One difference between the two promoters is that the *H2a-H2b* region lacks the CLAMP-binding GAGA repeats present in the *H3-H4* promoter (Rieder *et al*., 2017). Otherwise it is an intact, native bidirectional histone gene promoter. Consequently, the 12x^PR^ construct should retain the ability to initiate transcription from all RD histone genes. The 72nt size difference between the promoters provides a way to unambiguously distinguish between the 12x^PR^ and 12x^DWT^ array genotypes using PCR primers in the *H3* and *H4* coding regions in flies heterozygous (Fig. 1B) or homozygous (Fig. 1C) for the *HisC* deletion.

We first assessed whether the 12x^PR^ and 12x^DWT^ transgenes could support HLB formation in the presence of the endogenous histone genes in polytene chromosome spreads from 3^rd^ instar larval salivary glands. In these polyploid cells, the genome is amplified greater than 1000-fold and sister chromatids line up in register, resulting in large chromosomes providing high-resolution cytology. We visualized the HLB by immunofluorescence using antibodies that recognize one of several HLB components. These components include Multi sex combs (Mxc), the *Drosophila* ortholog of human NPAT, which is necessary for HLB assembly and histone gene expression (White *et al*., 2011; Terzo *et al*., 2015); histone pre-mRNA processing factors FLASH (Yang *et al*., 2009) and Lsm11, a component of the U7 snRNP (Godfrey *et al*., 2009; Burch *et al*., 2011); and Muscle wasted (Mute), an ortholog of human YARP (YY1AP-related protein 1) that is a putative repressor of histone gene expression (Bulchand *et al*., 2010). In these preparations, we also visualized both the endogenous histone locus and the ectopic, transgenic histone genes by FISH using a probe derived from the entire histone repeat unit. We observed HLB formation at the control 12x^DWT^ transgenic locus but not at the 12x^PR^ transgenic locus (Fig. 1D). These data are consistent with our previous results that sequences within the *H3-H4* promoter are necessary for HLB assembly in the presence of the endogenous histone genes (Salzler *et al*., 2013).

Both single copy transgenic histone gene repeats that fail to form an HLB (Salzler *et al*., 2013) and Mxc mutants that don’t form an HLB (Terzo *et al*., 2015) result in reduced histone mRNA levels, suggesting that the HLB is necessary for efficient histone mRNA biosynthesis. We therefore tested whether the 12x^PR^ transgene could support histone gene expression in the absence of HLB assembly. To determine expression from the ectopic 12x^DWT^ or 12x^PR^ genes in the presence of the endogenous RD histone genes, we randomly prime cDNA from total RNA and then amplify a fragment of each coding region containing the restriction enzyme site that is changed in the DWT construct. By digesting the PCR fragment with the appropriate restriction enzyme and resolving the fragments by gel electrophoresis we can determine the relative level of expression from the transgenic and endogenous RD histone genes. For example, RT-PCR of the *H2a* gene results in the same size amplification product from both the transgenic and endogenous genes, but only the product from the endogenous histone genes is sensitive to digestion with XhoI (Fig. 1E). When assayed in the presence of the endogenous genes, we found that *H2a* was expressed from the 12x^DWT^ at a higher level than from the 12x^PR^ transgene, even though identical promoters drove *H2a* in each transgenic array (Fig. 1E, Fig. 2A, lanes 2 and 4; Fig. 2B, lanes 2 and 3). These data are consistent with our previous observations using transgenes with a single histone gene repeat unit (Salzler *et al*., 2013; Terzo *et al*., 2015). Together these results demonstrate that the *H3-H4* promoter is required both for HLB formation and histone gene expression in the presence of the endogenous RD histone genes.

**Figure 2.**
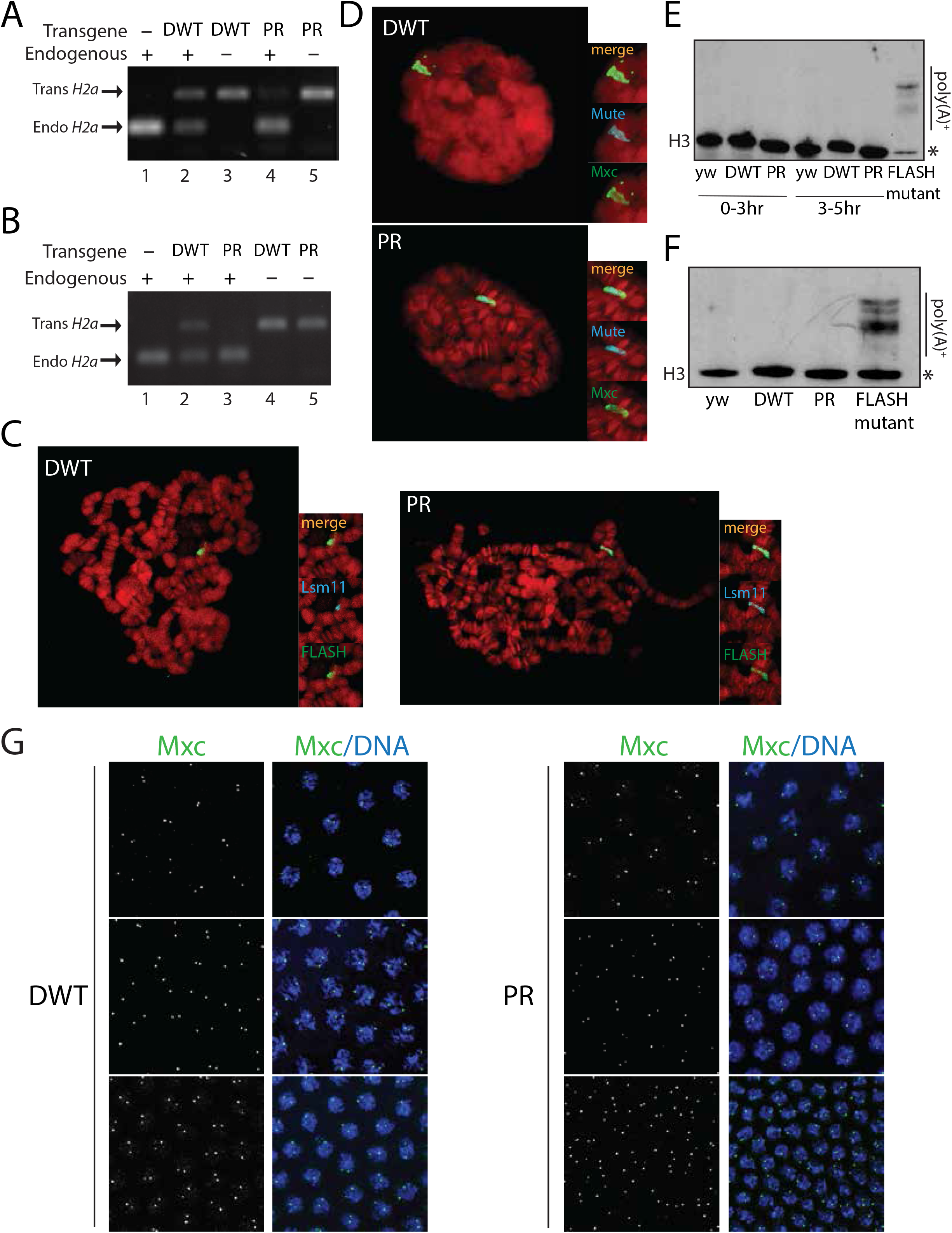
The PR transgene is expressed and forms an HLB in the absence of the endogenous RD histone genes. **A, B)** RNA was isolated from 5-7 hr embryos (A) or 3^rd^ instar larvae (B) containing the 12x^DWT^ or 12x^PR^ transgene in the presence or absence of *HisC*, with wild-type embryos as a control (lane 1). RT-PCR products using *H2a* primers were digested with *Xho I* and visualized using a 1.5% agarose gel (which does not resolve the two restriction fragments from the endogenous genes as in Fig. 1E). **C, D)** Polytene chromosomes from *HisC* deletion homozygous 3^rd^ instar larval salivary glands rescued by either the 12x^DWT^ or 12x^PR^ transgene stained with antibodies against HLB components Lsm11 and FLASH (C) or Mxc and Mute (D). **E, F)** Total RNA from 0-3h or 3-5h embryos (D) or 3^rd^ instar larvae (E) from wild-type (yw) or HisC deletion homozygotes containing either the 12x^DWT^ or 12x^PR^ transgene analyzed by Northern blotting using a radiolabeled *H3* coding region probe. * = normally processed H3 mRNA. A FLASH mutant that cannot recruit the processing machinery to the HLB (Tatomer *et al*., 2016b) was used as a positive control for production of poly(A)+ histone mRNA. **G)** Syncytial, embryos from *HisC* deletion homozygous stocks rescued by 12x^DWT^ or 12x^PR^ transgenes and stained with Mxc to monitor HLB formation in development. 3 consecutive syncytial nuclear cycles (top to bottom) were analyzed.

### A histone gene array lacking the H3-H4 promoter forms HLBs and is expressed in the absence of the endogenous genes

To assess the biological activity of the 12x^PR^ transgene, we determined the developmental outcome of having a 12x^PR^ transgene as the only zygotic source of histone mRNA. Due to large stores of maternal histone protein and mRNAs, embryos homozygous for a deletion that removes the endogenous RD histone gene array develop normally through S phase of cell cycle 14, but require zygotic RD histone gene expression for progression through S phase of cycle 15 and beyond (Gunesdogan *et al*., 2010, 2014). Consequently, embryos lacking RD histone genes arrest at nuclear cycle 15 and do not hatch. As noted above, this embryonic lethality is rescued by a single 12x^DWT^ transgene, which supports development of histone deletion progeny into viable, fertile, adult flies. Surprisingly, we found that embryos lacking endogenous histone genes and containing the 12x^PR^ transgene hatched and developed into nearly the expected number of fertile adults without any overt developmental delays, similar to embryos lacking endogenous histone genes and rescued by the 12x^DWT^ transgene. These data suggest that the 12x^PR^ transgene provides normal amounts of histone gene function when the endogenous RD histone genes are absent.

We interrogated this unexpected result more thoroughly by analyzing 1) histone gene expression, 2) HLB formation, and 3) histone pre-mRNA processing in both 5-7 hour old embryos and 3^rd^ instar larvae lacking endogenous histone genes and containing either the 12x^DWT^ or 12x^PR^ transgene. In both genotypes, all histone mRNA in either 5-7 hr embryos (Fig. 2A, lanes 3 and 5) or 3^rd^ instar larvae (Fig. 2B, lanes 4 and 5) was derived only from the ectopic histone gene array, indicating that the maternal histone mRNA stores had been degraded normally. We observed robust histone gene expression from the 12x^PR^ transgene when the endogenous histone genes are absent, in stark contrast with the low level of expression from the 12x^PR^ transgene when endogenous histone genes are present (Fig. 2A, lane 4; Fig. 2B lane 3).

High levels of histone gene expression are strongly correlated with the ability to form an HLB (Salzler *et al*., 2013; Terzo *et al*., 2015; Rieder *et al*., 2017). Given that the 12x^PR^ can support histone gene expression in the absence of the endogenous genes, we assayed for HLB formation in polytene chromosome spreads from 3^rd^ instar larval salivary glands and in embryos. We detected robust HLB formation at the 12x^PR^ transgenic locus with antibodies against FLASH and Lsm11 (Fig. 2C), or Mxc and Mute (Fig. 2D), similar to that observed at the 12x^DWT^ locus. Thus, the 12x^PR^ transgene, which lacks *H3-H4* promoter sequences, can support HLB formation and histone gene expression in the absence of endogenous RD histone genes.

### Histone mRNA from the 12x^PR^ transgenic locus is properly processed

An important function of the HLB is concentrating factors to promote efficient histone pre-mRNA processing (Tatomer *et al*., 2016b). Therefore, we reasoned it was possible that the attenuated 12x^PR^ gene array might affect other aspects of histone mRNA biosynthesis, including pre-mRNA processing. All *Drosophila* RD histone genes contain cryptic polyadenylation signals downstream of the normal processing sites that are only used when the histone pre-mRNA processing reaction is compromised, resulting in the production and accumulation of poly(A)+ histone mRNA (Lanzotti *et al*., 2002; Godfrey *et al*., 2009; Tatomer *et al*., 2016b). We examined pre-mRNA processing efficiency in flies rescued by the 12x^PR^ transgene. Our PCR based assays to detect histone mRNA expression cannot differentiate between properly processed and misprocessed histone mRNAs. We therefore used northern blotting with a probe against the *H3* coding region to determine whether histone pre-mRNA was efficiently processed. In contrast to a *FLASH* mutant, which expresses large amounts of poly(A)+ histone mRNA, we did not detect poly(A)+ histone mRNA from early embryos (Fig. 2E) or from whole 3^rd^ instar larvae (Fig. 2F) in *HisC* deletion animals rescued by the 12x^PR^ transgene. Note also that the levels of histone mRNA from the 12x^PR^ transgenes are similar to wild type levels. These results indicate that the HLB formed on the 12x^PR^ array in the absence of endogenous histone genes supports efficient histone pre-mRNA processing.

### HLB assembly at the 12x^PR^ transgenic locus occurs at the normal time during embryogenesis

The above data suggest that the 12x^PR^ represents an attenuated histone gene array that provides normal biological function when not in competition with the wild type endogenous histone genes. To further explore this model, we determined if an HLB assembles on the 12x^PR^ array in the early embryo at the same time that it assembles on the 12x^DWT^ array. The HLB begins assembling in syncytial blastoderm embryos just prior to the onset of zygotic histone transcription (White *et al*., 2007; White *et al*., 2011). We previously reported that a “proto-HLB” consisting of Mxc and FLASH forms in nuclear cycle 10, followed by the onset of zygotic histone gene expression and further recruitment of additional HLB components (Mute and Lsm11) in cycle 11 (White *et al*., 2011; Salzler *et al*., 2013). To determine if the HLB forms at the 12x^PR^ transgenic locus with normal timing, we stained syncytial blastoderm embryos lacking endogenous histone genes and rescued by a 12x^PR^ transgene with antibodies against Mxc (Fig. 2G). In these experiments, HLB assembly during the syncytial blastoderm cycles was indistinguishable from that of histone deletion embryos rescued by the control 12x^DWT^ transgene. Thus, in the absence of the endogenous genes, an HLB assembles on the 12x^PR^ array at the same time in early development as it does on a wild type array.

### CLAMP is present in the 12x^PR^ HLB in the absence of the endogenous RD histone genes

The *H3-H4* promoter is highly conserved among *Drosophilids* and contains conserved GAGA repeats (Salzler *et al*., 2013; Rieder *et al*., 2017), which we previously showed are essential for HLB formation and expression of RD histone genes in the presence of the endogenous histone locus (Rieder *et al*., 2017). Although in vitro these repeats can bind both the zinc-finger GA-repeat binding protein CLAMP, and the major *Drosophila* GAGA repeat binding protein GAF (GAGA factor*; trithorax-like*) (Gilmour *et al*., 1989) only CLAMP is bound to the histone locus in wild-type animals (Rieder *et al*., 2017). The *H2a-H2b* promoter and the rest of the histone repeat unit do not contain any GAGA repeats longer than 4 nts, and CLAMP preferentially binds to longer GAGA repeats (Kuzu *et al*., 2016). We asked whether CLAMP is recruited to the 12x^PR^ transgene in salivary gland polytene chromosomes. As with all other HLB components we tested, CLAMP is not recruited to the 12x^PR^ transgene or a 12x histone gene array in which the GAGA sequences are replaced with *lacO* binding sites (GA^M^, Fig. 3A) when the endogenous RD histone genes are present (Rieder *et al*., 2017). Surprisingly, we found that in the absence of endogenous RD histone genes, CLAMP (Fig. 3C), but not GAF (Fig. 3E), is recruited to the 12x^PR^ transgenic locus with similar intensity to CLAMP recruitment to the 12x^DWT^ transgenic locus (Fig. 3B, D). Furthermore, in this genotype the 12x GA^M^ array supports *H2a* gene expression in the absence of the endogenous genes (Fig. 3F). These data indicate that CLAMP can be recruited to a histone gene array lacking GAGA repeats when the preferred GAGA binding sites within the *H3-H4* promoter at the endogenous RD histone locus are absent. The *H3* and *H4* genes are transcribed, indicating that CLAMP is not an essential DNA-binding transcription factor for the *H3* and *H4* genes, but likely serves as a factor that alters chromatin structure (Rieder et al, 2017).

**Figure 3.**
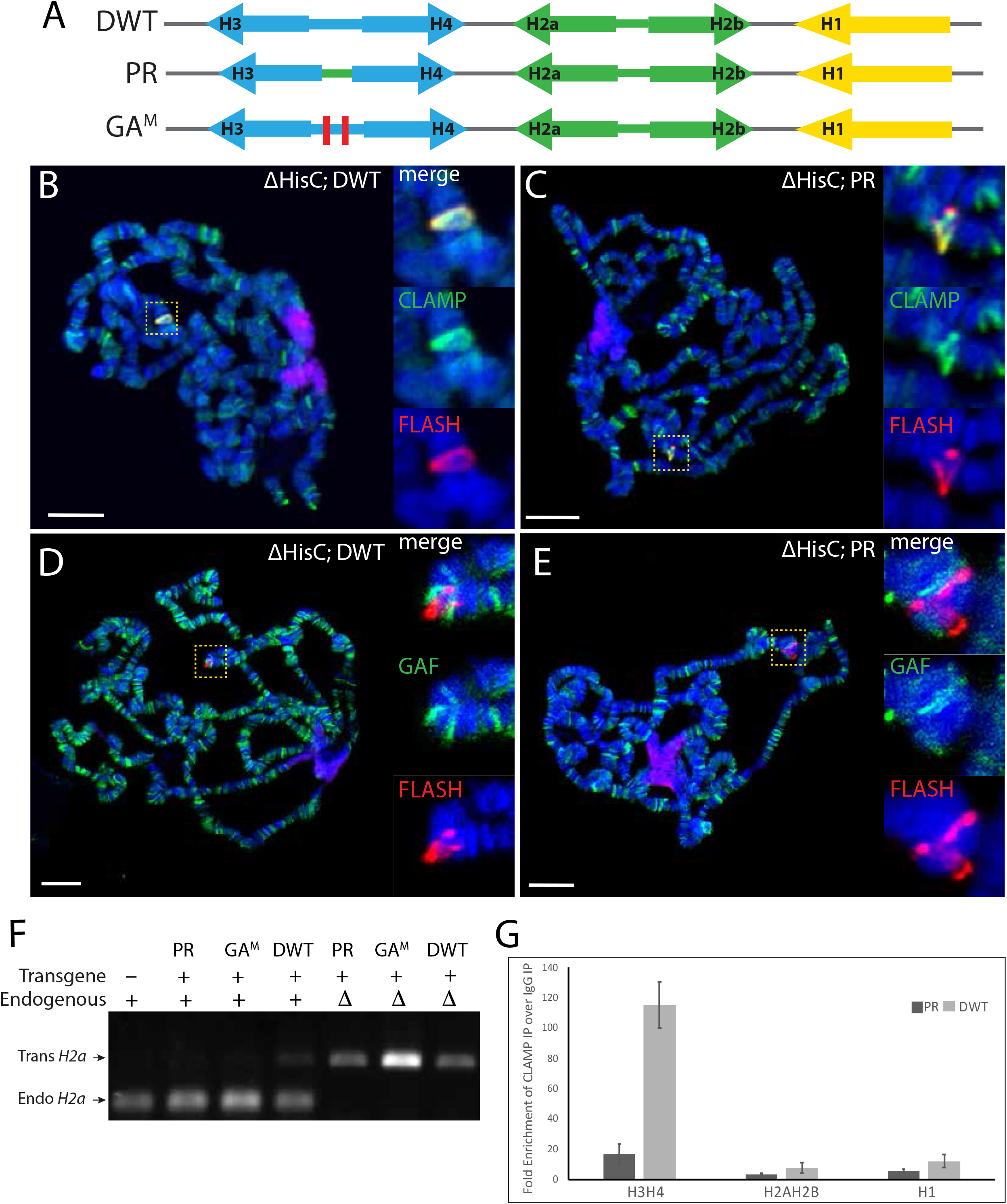
CLAMP but not GAF is recruited to HLBs that form in the absence of GA repeats. **A)** Schematic of the WT, PR, and GA mutant (GA^M^) synthetic histone repeats. For GA^M^, GA sequences in the *H3-H4* promoter were mutated to *lacO* sites or scrambled (Rieder *et al*., 2017). **B-E)** Polytene chromosome squashes from 3^rd^ instar larval salivary glands lacking endogenous histone genes and rescued by a either a 12x^DWT^ transgene (B,D) or a 12x^PR^ transgene (C,E) were stained with antibodies against FLASH and CLAMP (B,C) or FLASH and GAF (D,E). Scale bar 10 microns. **F)** RT-PCR analysis with *H2a* primers performed on cDNA from 5-7 hr embryos of indicated genotypes and visualized with ethidium bromide on a 0.8% agarose gel. **G)** 2-4 hr old *HisC* homozygous deletion embryos rescued by a 12x^DWT^ or 12x^PR^ transgene were analyzed by ChIP-PCR using primers recognizing the indicated promoters.

We carried out ChIP-qPCR experiments on embryos containing only the 12x^DWT^ array or the 12x^PR^ array to determine whether CLAMP is interacting with either the *H2a-H2b* or *H3-H4* genes. In agreement with ChIP-seq results on the endogenous genes, which showed that CLAMP is localized precisely to the *H3-H4* promoter (Rieder *et al*., 2017), CLAMP was bound to the *H3-H4* promoter but not to the *H2a-H2b* genes or promoter on the 12x^DWT^ array (Fig. 3G). In contrast, CLAMP was not bound to either gene pair on the 12x^PR^ array (Fig. 3G), despite our observation that CLAMP is present in the HLB at the 12x^PR^ array (Fig. 3C). These data suggest that CLAMP can be recruited to RD histone genes by protein-protein interactions independently of its binding to DNA.

### A wild type array can activate the 12x^PR^ array when present in trans at the homologous locus

The results above demonstrate that the 12x^PR^ transgenic locus does not form an HLB in the presence of the endogenous histone genes unlike the wild type 12x^DWT^ transgene, which does. We tested whether juxtaposing a wild type histone gene array near the 12x^PR^ locus would nucleate a functional HLB and activate expression from the 12x^PR^ transgene. Our BAC based transgenes are inserted into the genome via site-specific recombination at the same chromosomal location. We created flies in which the 12x^PR^ was placed at position VK33 on chromosome 3 in trans to the 12x^HWT^ transgene used in our initial studies (McKay *et al*., 2015), and analyzed these transgenes in the presence of the endogenous RD histone genes on chromosome 2. We examined HLB formation at the ectopic VK33 location by staining whole mount salivary glands with antibodies against Mxc and FLASH. Using this approach, we detect only a single, large HLB on the endogenous RD histone genes in wild type nuclei (Fig. 4A-1, 4B). When 12x^PR^ was the only transgene present in addition to the endogenous RD histone genes, none of the nuclei contained an ectopic HLB and we only detected the endogenous HLB (Fig. 4A-2, 4B), consistent with the results of staining spread salivary gland polytene chromosomes (Fig. 1D). In contrast, we observed formation of a second small HLB in addition to the single large endogenous HLB in 100% of the 12x^HWT^/12x^PR^ nuclei (Fig. 4A-5, 4B), as expected since this HLB also forms when only the 12x^HWT^ array is present at this site.

**Figure 4.**
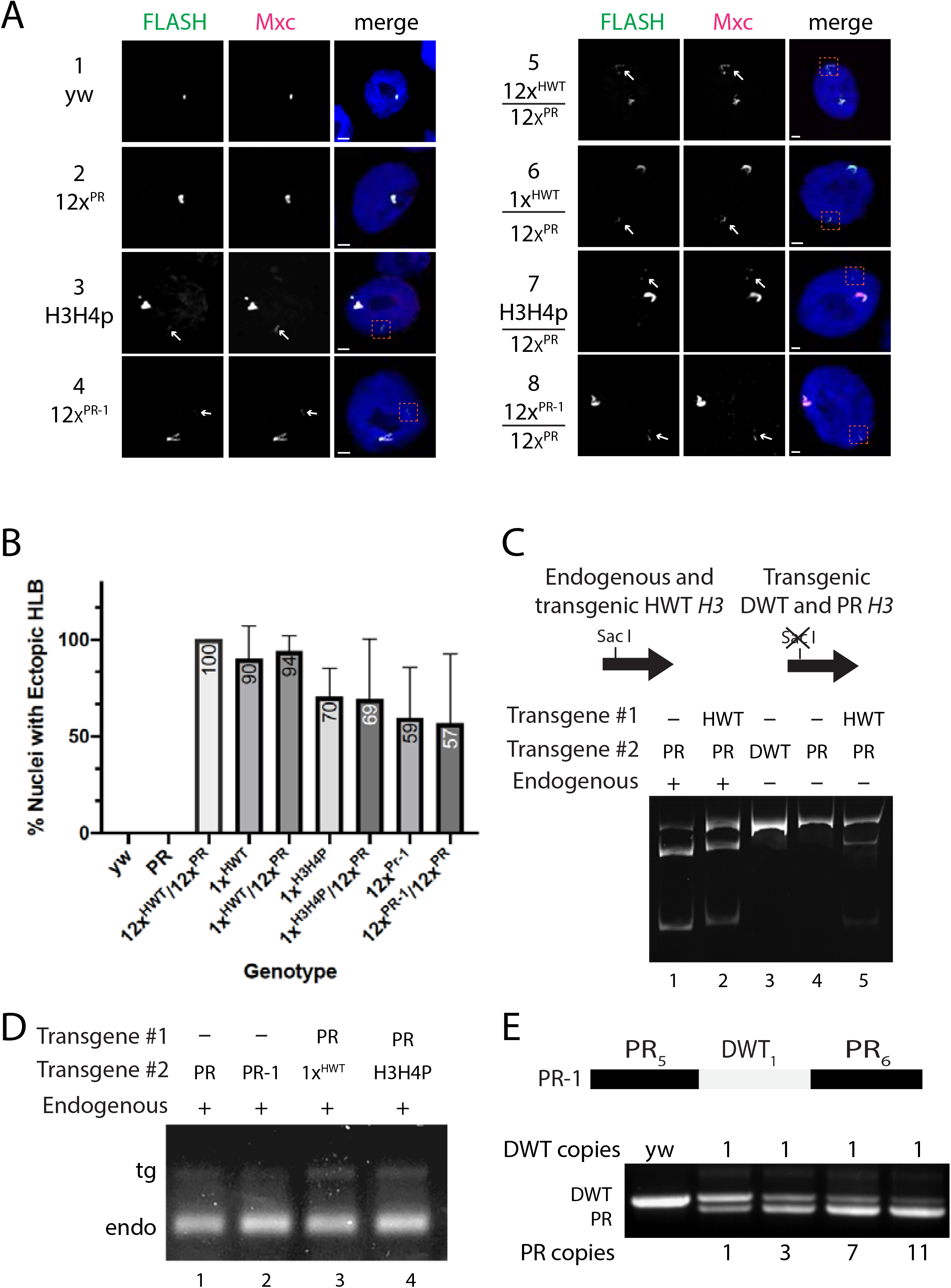
Activation of PR transgenes by wild type histone gene arrays. A-1 to A-8) Intact salivary gland nuclei from 3^rd^ instar larvae of the indicated genotypes, all of which contained *HisC* and stained for the HLB components Mxc and FLASH. Ectopic HLBs are indicated by arrows and dashed boxes, and were present in all genotypes except the 12x^PR^. Scale bar 10 microns. B) Quantification of the percentage of ectopic HLBs in the salivary gland nuclei in panel B. C) Schematic of the assay used to distinguish 12x^PR^ from 12x^HWT^ cDNA. RT-PCR amplification of *H3* mRNA from 5-7 hr old embryos of the indicated genotypes, followed by digestion with SacI and visualized on an 8% polyacrylamide gel. D) RT-PCR amplification of *H2a* mRNA from larvae of indicated genotypes, followed by digestion with XhoI and visualization on an agarose gel. E) Top: Schematic of the BAC-based PR-1 histone array containing 11 PR and 1 DWT repeat unit. Bottom: Analysis of BAC constructs containing WT and PR repeats using PCR as in panel 1B and C.

We measured histone gene expression from the 12x^PR^ transgene in these genotypes. In the 12x^HWT^ array the histone *H2a* gene, and not the other histone genes, is marked with a restriction site change (McKay *et al*., 2015), whereas the 12x^PR^ array has all five histone genes marked with a restriction site change. Therefore, to specifically detect histone gene expression from the 12x^PR^ transgene, we digested embryonic *H3* RT-PCR products with SacI, whose recognition site is missing from the 12x^PR^ *H3* gene but is present in both the endogenous and 12x^HWT^ *H3* genes (Fig. 4C). Strikingly, the 12x^PR^ transgene supports stronger *H3* gene expression in the presence compared to the absence of the endogenous histone genes when present in trans with the 12x^HWT^ transgene (Fig. 4C, lanes 1 and 2). These data demonstrate that the wild type histone sequences in the 12x^HWT^ were able to activate the 12x^PR^ transgenic locus in the presence of the endogenous histone genes, likely by nucleating ectopic HLB formation that encompasses the paired homologous chromosomes. This result is similar to transvection, a phenomenon in which transcription of a gene can be activated by an enhancer located in trans through pairing of homologous chromosomes (Duncan, 2002; Fukaya and Levine, 2017).

### Transgenes with a single copy of the wild type repeat partially activate a 12x array in trans

Since 12 copies of a wild type histone repeat at the same integration site stimulated expression from the 12x^PR^ transgene, we tested whether a single copy could do the same using two different approaches; one where the single copy is present *in trans* and the other where it is present in *cis*. For the first approach, we inserted a 1x HWT transgene at the VK33 integration site. In these strains we detected the HLB in 90% of the salivary gland nuclei in the presence of the endogenous genes (Fig. 4B), similar to what we observed previously (Salzler *et al*., 2013). When this 1x HWT transgene was placed *in trans* with the 12x^PR^ array, we detected an ectopic HLB in 94% of the nuclei examined (Fig. 4A-6, 4B). We tested whether there was any activation of expression of the 12x^PR^ array using our RT-PCR assay. There was a modest activation of the expression 12x^PR^ array, but much weaker than when the 12x^HWT^ was *in trans* with the 12x^PR^ as measured by the ratio of the bands (Fig. 4D, lanes 1 and 3).

We next tested if including a single wild type repeat in the 12x^PR^ array could stimulate HLB formation and histone mRNA expression from that array in the presence of the endogenous genes. We created a 12x array (12x^PR-1^) containing one wild type histone repeat unit in the center of 11 PR repeat units (Fig. 4E). Like 12x^PR^, the 12x^PR-1^ transgene rescued a deletion of the endogenous histone genes, resulting in viable and fertile adults and indicating that it is likely fully active when present as the only source of RD histone genes. We then examined HLB formation in intact salivary glands from animals containing both the 12x^PR-1^ transgene and the endogenous RD histone genes. In this genotype, we detected HLB formation at the ectopic 12x^PR-1^ transgenic locus in 59% of the nuclei (Fig. 4A-4, 4B), compared to 0% of nuclei from the 12x^PR^. We also analyzed expression from this array in the presence of the endogenous RD histone genes and observed only modest activation of histone mRNA expression (Fig. 4D, lane and less than the level of expression when the 1X WT was *in trans* to the 12X array (Fig. 4D, lane 3). Thus, a single histone repeat unit can stimulate HLB formation, but activates expression of neighboring genes better when placed *in trans* (i.e. on the other homolog) rather than *in cis* (i.e. in the same 12x array).

We next tested a transgene containing only the *H3-H4* promoter with no coding or 3’ end sequences, which we previously demonstrated could form an HLB in salivary glands when inserted into a ectopic site (Salzler *et al*., 2013). Ectopic HLBs were detected in 70% of the salivary glands when only the *H3-H4p* promoter transgene was present and 69% when it was placed *in trans* the 12x^PR^ array (Fig. 4A-3, 4A-7, 4B). We assayed the expression of the *H3* gene from the 12x^PR^ when the *H3-H4p* transgene was in trans to the 12x^PR^ array. We detected low level expression, similar to the expression detected when the 1xWT transgene was present opposite the 12x^PR^ array (Fig. 4D, lane 4). Collectively, these data demonstrate that the presence of an HLB nucleating sequence *in trans* to 12x^PR^ can induce formation of an ectopic HLB and histone gene expression in the presence of the endogenous histone genes. Further, these data emphasize that the *H3-H4* promoter is a critical element in promoting HLB formation, consistent with our previous observations (Salzler *et al*., 2013; Rieder *et al*., 2017).

## Discussion

In this study, we used our histone gene replacement platform to analyze the cis acting elements within the *Drosophila* histone repeat unit that are necessary for HLB formation and histone gene expression. Previously we showed using a single, transgenic histone gene repeat unit that the promoter region of the divergently transcribed *H3-H4* gene pair is capable of stimulating HLB formation (Salzler *et al*., 2013). We subsequently further mapped this functionality using a 12x gene array to conserved GAGA repeats in this region that are targeted by the CLAMP protein (Rieder *et al*., 2017). Here, we present evidence that although a 12x^PR^ histone gene array devoid of the *H3-H4* promoter and lacking CLAMP binding elements cannot assemble an HLB in the presence of the ∼100 RD histone gene copies at the endogenous locus (*HisC*), it surprisingly can rescue homozygous deletion of *HisC* and fully support the entire *Drosophila* life cycle. In the *HisC* deletion background, the 12x^PR^ array assembles an HLB and expresses the same amount of properly processed histone mRNAs as the endogenous genes or as a 12x^DWT^ wild type array. Below we discuss the implications of these observations on HLB assembly and organization.

### The RD histone locus stimulates HLB formation in Drosophila

Biomolecular condensates form via a seeding event that promotes a high concentration of factors at a discrete location, leading to recruitment of additional factors that ultimately results in a structure that can be observed by light microscopy (Gomes and Shorter, 2019). A number of putative seeding events for biomolecular condensates have been described (Dellaire *et al*., 2006; Dundr, 2011; Mao *et al*., 2011; Shevtsov and Dundr, 2011; White *et al*., 2011), but in many cases the precise mechanism of seeding is not known. Nucleic acids, particularly RNA, have been proposed to seed different nuclear bodies. Both the nucleolus and the HLB are associated with specific genomic loci (Mao *et al*., 2011; Shevtsov and Dundr, 2011; Salzler *et al*., 2013), and it is likely that the DNA (or chromatin) and/or nascent RNA at the locus participates in the seeding event. The activation of zygotic transcription of rRNA leads to the precise spatiotemporal formation of the nucleolus in *Drosophila* embryos (Falahati *et al*., 2016). In the absence of rDNA, *Drosophila* nucleolar components still form high concentration assemblies, but these are smaller, more numerous, and do not form at the same time in the early embryo as the wild type nucleolus.

*Drosophila* HLB components also stochastically assemble smaller and more unstable foci in embryos lacking the RD histone locus (White *et al*., 2007; Salzler *et al*., 2013; Hur, 2019), suggesting that HLBs and the nucleolus form similar seeding events. Indeed, the dynamics of HLB assembly in single early *Drosophila* embryos display properties consistent with liquid-liquid phase transition seeded by *HisC* (Hur, 2019). Blocking transcription in the early embryo prevents normal HLB growth (Hur, 2019), and a defective *H3-H4* promoter (with mutated TATA boxes) does not support HLB formation in the context of a single copy histone gene repeat in salivary glands (Salzler *et al*., 2013). These data suggest that active transcription is essential for forming a complete HLB.

It is important to note that HLBs assemble and persist in non-proliferating *Drosophila* tissues that do not express histone mRNAs (Liu *et al*., 2006; White *et al*., 2007), and are also present in G0/G1 mammalian cells (Ma *et al*., 2000; Zhao *et al*., 2000). Histone gene expression is activated as a result of phosphorylation of Mxc/NPAT by Cyclin E/Cdk2, resulting in changes in the HLB that promote histone gene transcription and pre-mRNA processing (Wei *et al*., 2003; Ye *et al*., 2003; White *et al*., 2011). We propose that in early embryonic development the histone locus DNA and/or chromatin seeds HLB assembly in *Drosophila*, with the *H3-H4* promoter region being particularly important. We further propose that subsequent transcriptional activation of histone genes then drives HLB growth and maturation.

Formation of an HLB on a transgenic RD histone gene array requires that this array compete effectively with the endogenous *HisC* locus for recruitment of HLB components. This is the situation with 12x^HWT^ and 12X^DWT^ arrays, which form HLBs in the presence of *HisC*. These results also indicate that there are no other elements within *HisC* that are necessary for HLB formation. Because the 12x^PR^ array does not form an HLB in the presence of *HisC*, but it does so in the absence of *HisC*, we hypothesize that the endogenous RD histone gene array sequesters critical HLB components, likely including Mxc and CLAMP, thereby preventing HLB assembly at the transgenic locus. By removing the *H3-H4* promoter from the transgene, we removed an element that provided additional interactions with HLB components, notably CLAMP, weakening the overall ability of the locus to stably nucleate an HLB.

### HLB assembly involves multiple interactions among components

Interactions among multivalent proteins, or multivalent protein-nucleic acid interactions, are driving forces in the assembly of biomolecular condensates (Shin and Brangwynne, 2017). Mxc is likely the critical factor that together with histone genes seeds *Drosophila* HLB formation and activates histone gene expression. Mxc is a large (∼1800 aa) protein that oligomerizes in vivo and likely provides a scaffold for multivalent interactions (Terzo *et al*., 2015). A C-terminal truncation mutant of Mxc that fails to recruit histone pre-mRNA processing factors still forms an HLB and activates histone gene expression at sufficient levels to complete development, underscoring the multivalent nature of Mxc (White *et al*., 2011; Landais *et al*., 2014; Tatomer *et al*., 2016b).

Surprisingly, the HLB that assembles on the 12x^PR^ array in the absence of *HisC* contains CLAMP, even though we have removed all of the known CLAMP binding sites from the histone repeat. Although CLAMP may bind another sequence in the 12x^PR^ array, no other favorable GAGA repeats are present, and we were unable to detect CLAMP bound to any of the histone promoters by ChIP-qPCR experiments. <u>More likely,</u> CLAMP may interact with other HLB components, possibly Mxc or the Mxc-FLASH complex, providing multivalent contacts between CLAMP and other HLB components. Deleting the GAGA sequences from the *H3-H4* promoter did not affect transcription of the *H3* or *H4* genes in the absence of *HisC*, suggesting that CLAMP’s major function is to promote HLB assembly and not to act as a DNA binding transcription factor. Supporting this interpretation is the observation that another, more abundant transcription factor that binds to GAGA repeats, GAF, is not found at the HLB unless CLAMP is absent (Rieder *et al*., 2017), consistent with CLAMP’s critical interactions with both the GAGA repeats and HLB factors in seeding the HLB.

Because the 12x^PR^ array is capable of assembling a completely functional HLB in the absence of *HisC*, the *H3-H4* promoter is not absolutely essential for HLB formation. One possibility is that there are multiple pathways for assembling functional HLBs. Previous work suggests that not all seeding events are equivalent in their ability to assemble biomolecular condensates. In artificial systems, changes in scaffold stoichiometry, which can stem from changes in valence, alter the recruitment of components (Banani *et al*., 2016). Further, mathematical modeling has revealed that scaffolds can nucleate distinct complexes when at different concentrations, and that this can qualitatively alter the transcriptional output (Yang and Hlavacek, 2011). Additionally, P-bodies can form in multiple ways through different protein-protein or protein-nucleic acid interactions, with different interactions predominating under different conditions (Rao and Parker, 2017). Therefore, different nucleators of the HLB (i.e. the *H3-H4* promoter or other sequences in the locus) may result in similar but not identical outcomes. Collectively our results suggest that HLB formation results from the contribution of many molecular interactions, and the loss of any single one may be overcome by other multivalent interactions within the body.

## Methods

### Culture condition and fly strains

Original fly strains and crosses were used as in McKay et al. 2015. Stocks were maintained on standard corn medium. Viability studies were performed as in (Penke *et al*., 2016).

### Transgenic histone gene array construction

Construction of the 5kb histone repeat designed for this study was performed using the HiFi DNA Assembly system from New England Biolabs. PCR amplification of fragments from existing histone repeats, in addition to in vitro synthesized gblocks (IDT) containing the desired nucleotide changes were employed for the building blocks of the reaction. Manufactures protocol was followed with slight modification to the incubation time of the reaction. Engineered 5kb histone repeats were then arrayed to 12 copies in pMultiBac as in (McKay *et al*., 2015). All histone gene arrays were inserted into the VK33 attP site on chromosome 3L (65B2) (Venken *et al*., 2006).

### Northern Analysis

Northern analysis was performed using a 7 M 6% urea acrylamide gel to resolve histone mRNAs. ∼1ug of RNA from embryos or larvae was used with a radio-labeled probe to the coding region of H3 as in (Lanzotti *et al*., 2002).

### Histone Expression Analysis

Total RNA was prepared in Trizol and cDNA synthesized with random hexamers using Superscript II (Invitrogen), according to the manufacturer’s instructions. RT-RCR was performed using gene-specific primers to H2a (McKay et al. 2015) and H3. Each reaction was performed at least 3 times with similar results. PCR products were digested using XhoI (H2a) or SacI (H3). Digested amplicons were run on an 8% acrylamide gel or a 1.5% agarose gel.

### Immunofluorescence

We used primary antibodies at the following concentrations: rabbit anti-CLAMP (1:1000), guinea pig anti-Mxc (1:2000), guinea pig anti-Mute (1:5000), rabbit anti-FLASH (1:2000), rabbit anti-Lsm10 (1:1000), mouse anti-MPM-2 (1:100; Millipore), rabbit anti-GAF (1:1000), mouse anti-LacI (1:1000; Millipore). We used Alexa fluor secondary antibodies (Thermo Fisher Scientific) at a concentration of 1:1000. In situ probes were detected using 15 µg/mL streptavidin-DyLight-488 (Vector Laboratories). Salivary gland dissections and squashes were performed as in (Tatomer *et al*., 2016b). Images were acquired with using a Zeiss Lsm710 with ZEN DUO software. Images were analyzed using ImageJ. Ectopic HLBs in embryos were quantified as (Salzler *et al*., 2013).

### Fluorescent in situ hybridization

FISH probes were made from a PCR product that spanned the entire wild-type histone repeat (primers [AAAGGAGGTTGGTAGGCAGC] and [ACGCTAGCGCTTTATCTGCA]). Biotinylated FISH probes were made with nick translation by incubating 1 µg of purified PCR product for 2 h at 15°C in a total of 50 µL containing 1×DNAPolI buffer (Fisher Optizyme); 0.05mM each of dCTP, dATP, and dGTP; 0.05 mM biotin-11-dUTP (Thermo Scientific); 10 mM 2-mercaptoethanol; 0.004 U of DNaseI (Fisher Optizyme); and 10 U of DNAPol I (Fisher Optizyme). The reaction was purified on a PCR purification column (Thermo Scientific) and diluted in hybridization buffer (2xSSC, 10%dextran sulfate, 50%formamide, 0.8 mg/mL salmon sperm DNA) to a final volume of 220 µL. FISH probes were diluted in hybridization mixture and added to slides containing salivary gland polytene chromosome spreads before heating. We added a coverslip, sealed it with rubber cement, and heated the slide for 2 min on a 91°C heat block. Slides were placed in a humid box and incubated at 37°C overnight. Immunostaining was then performed by incubating the slides in primary antibody overnight at 4°C in a humid box.

### Chromatin Immunoprecipitation

ChIP was performed according to (Blythe and Wieschaus, 2015) with the following particulars. 2-4 hours old embryos (∼200 embryos for each of three biological replicates) from *HisC/HisC*; 12x^DWT^ or 12x^PR^ strains were collected, fixed and stored at −80°C until further processing. Immunoprecipitation of chromatin preparations was performed using rabbit anti-CLAMP antibody (Rieder *et al*., 2017) and a polyclonal rabbit IgG (Millipore-Sigma, 12-370). qPCR was performed as described in (Urban *et al*., 2017). Three biological replicates and two technical replicates were used for each genotype. We normalized CLAMP IP target abundance against IgG IP target abundance. We analyzed data using a Student’s t-test, comparing target abundance between the DWT and PR lines.

## Acknowledgements

We thank Erica Larschan’s lab at Brown University for assistance with ChIP experiments. This work was supported by NIH grant R01GM058921 to R.J.D and W.F.M and F32GM109663, K99HD092625, and R00HD092625 to L.E.R.

---

**Supplemental Figure 1.**
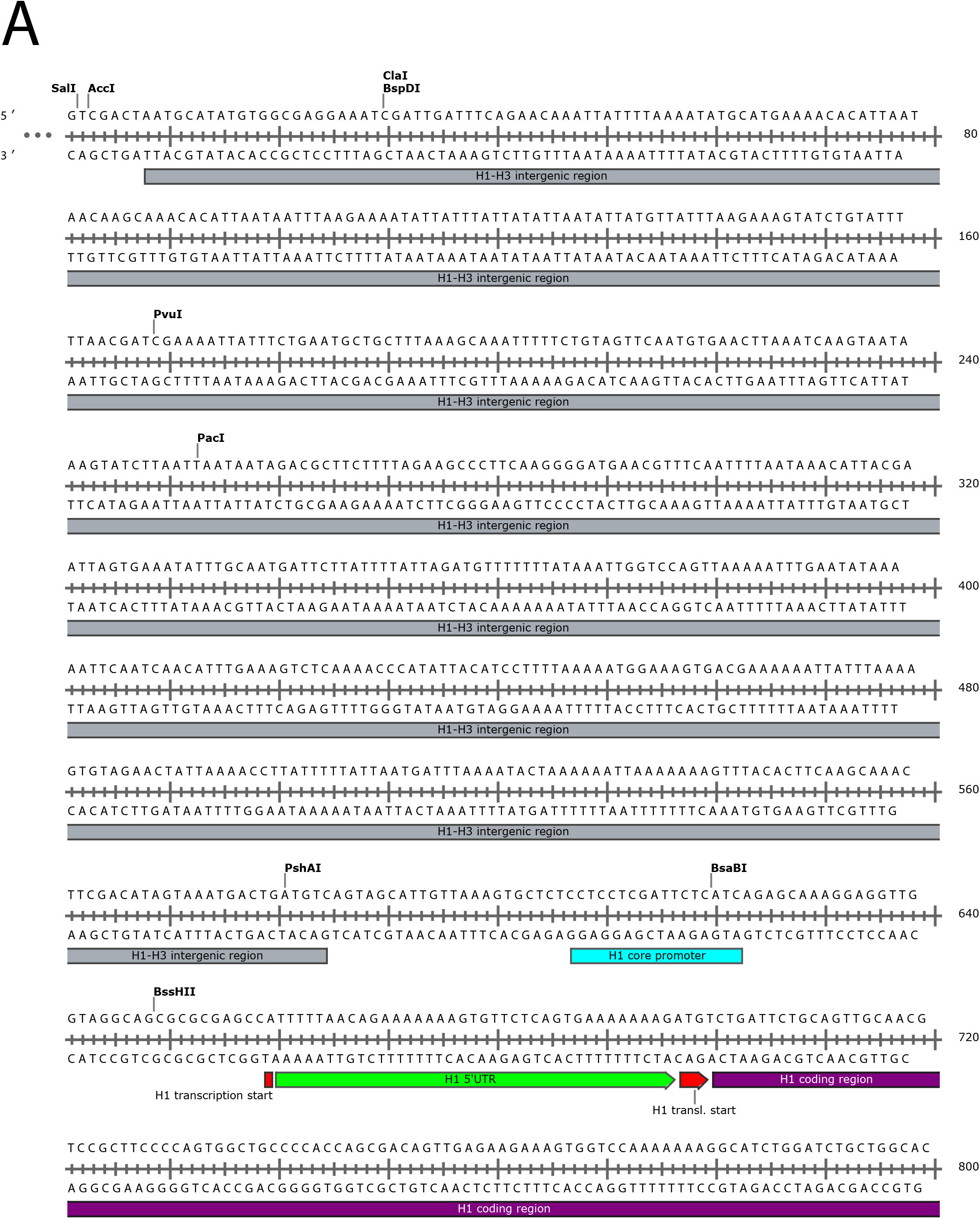
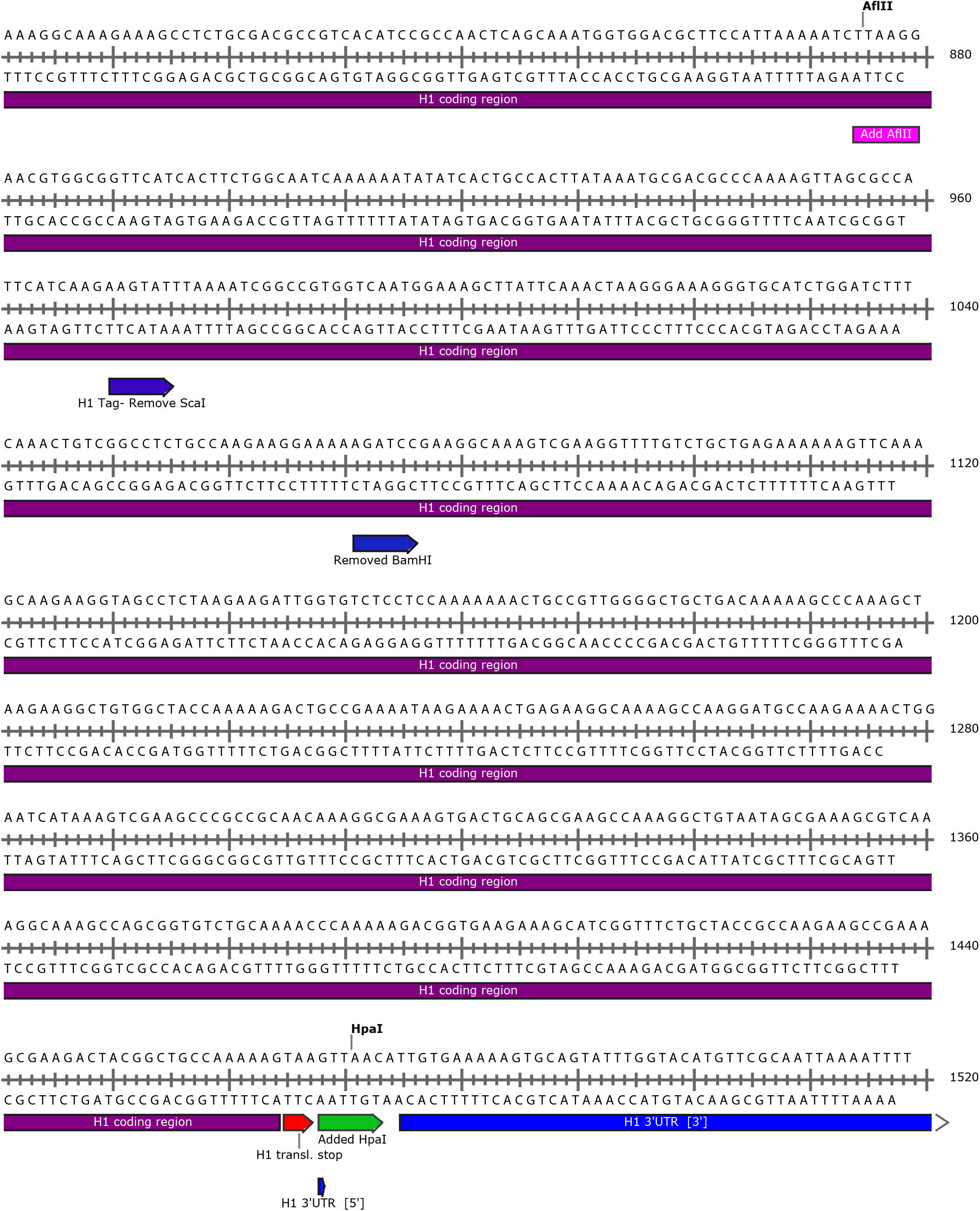
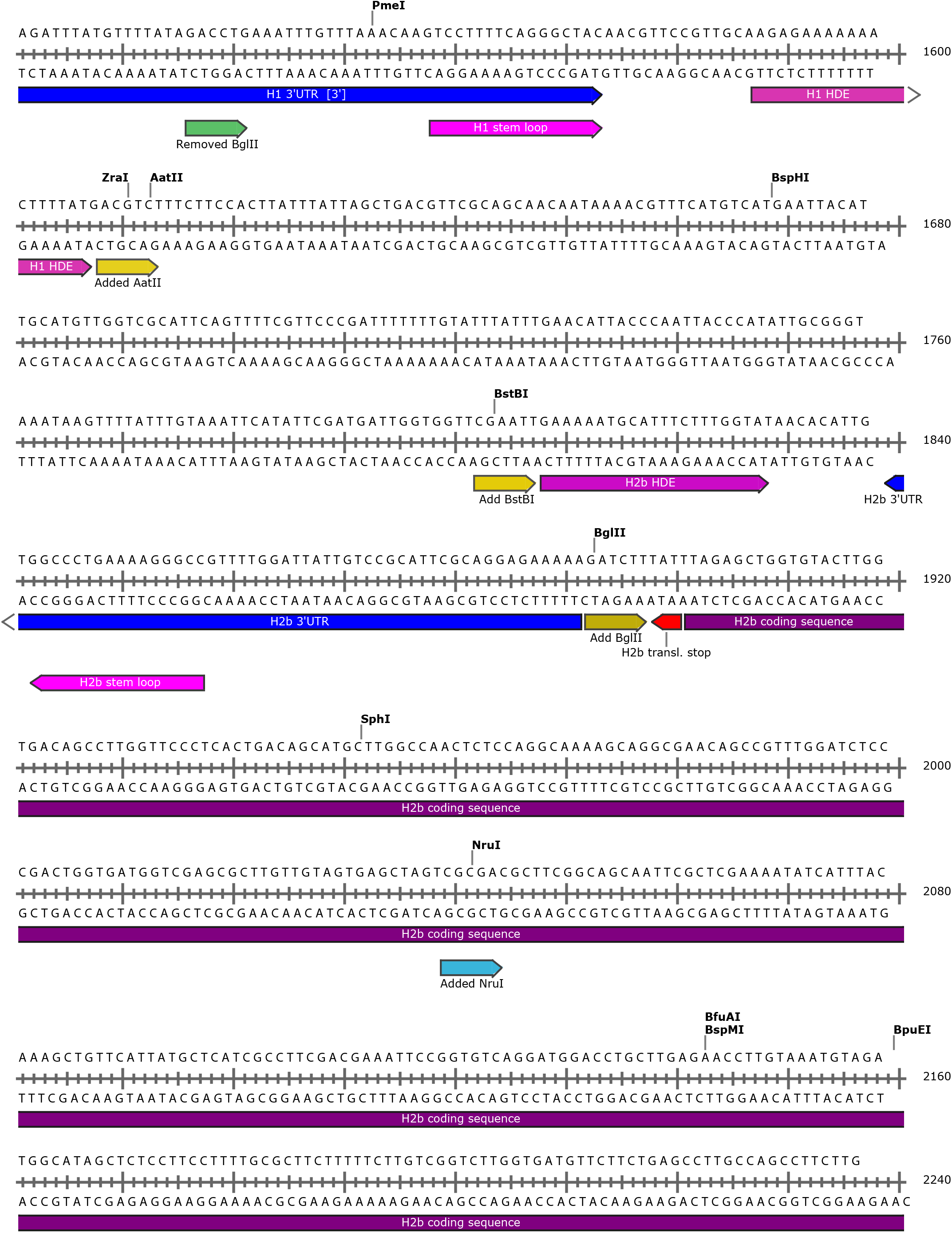
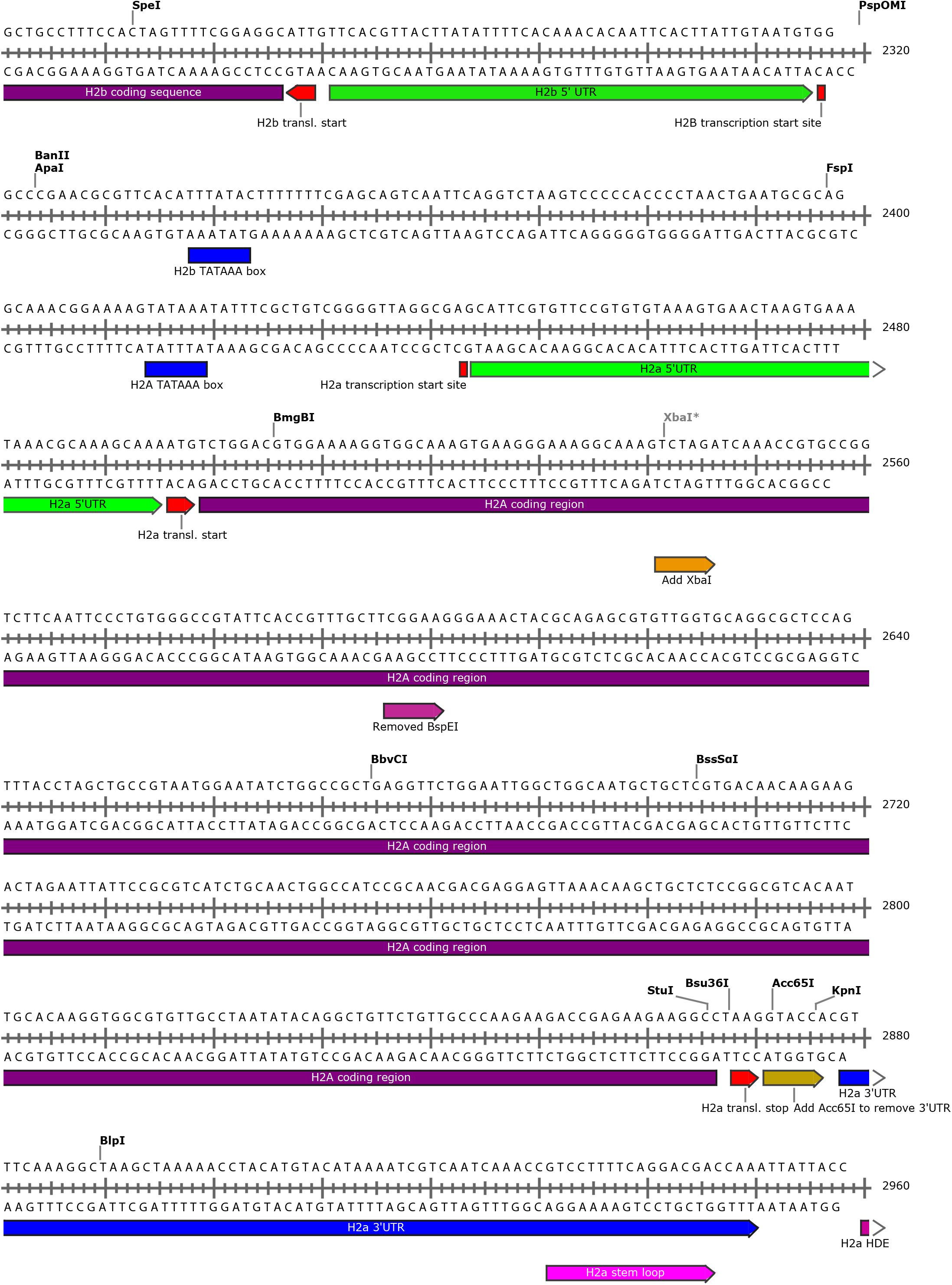
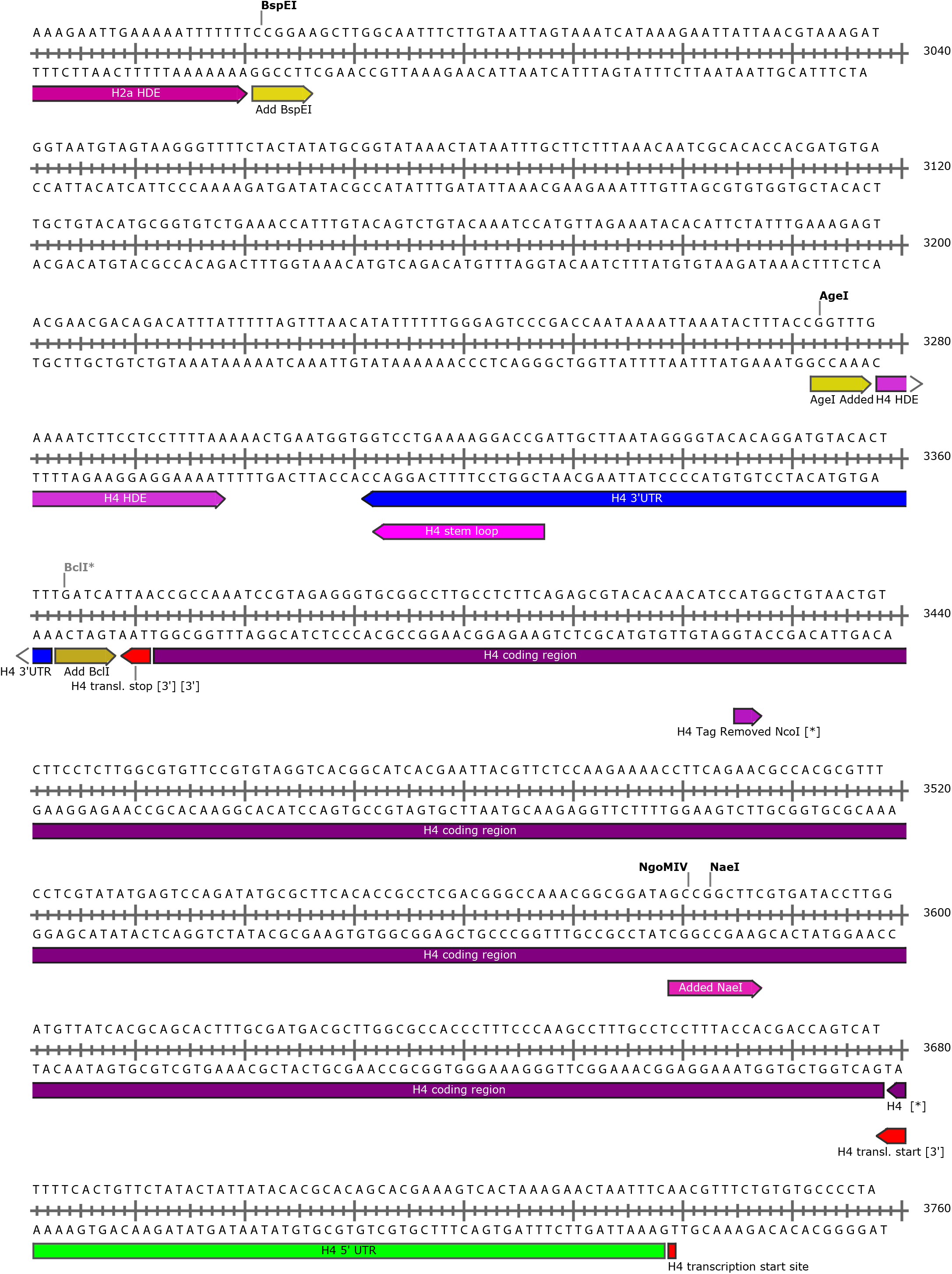
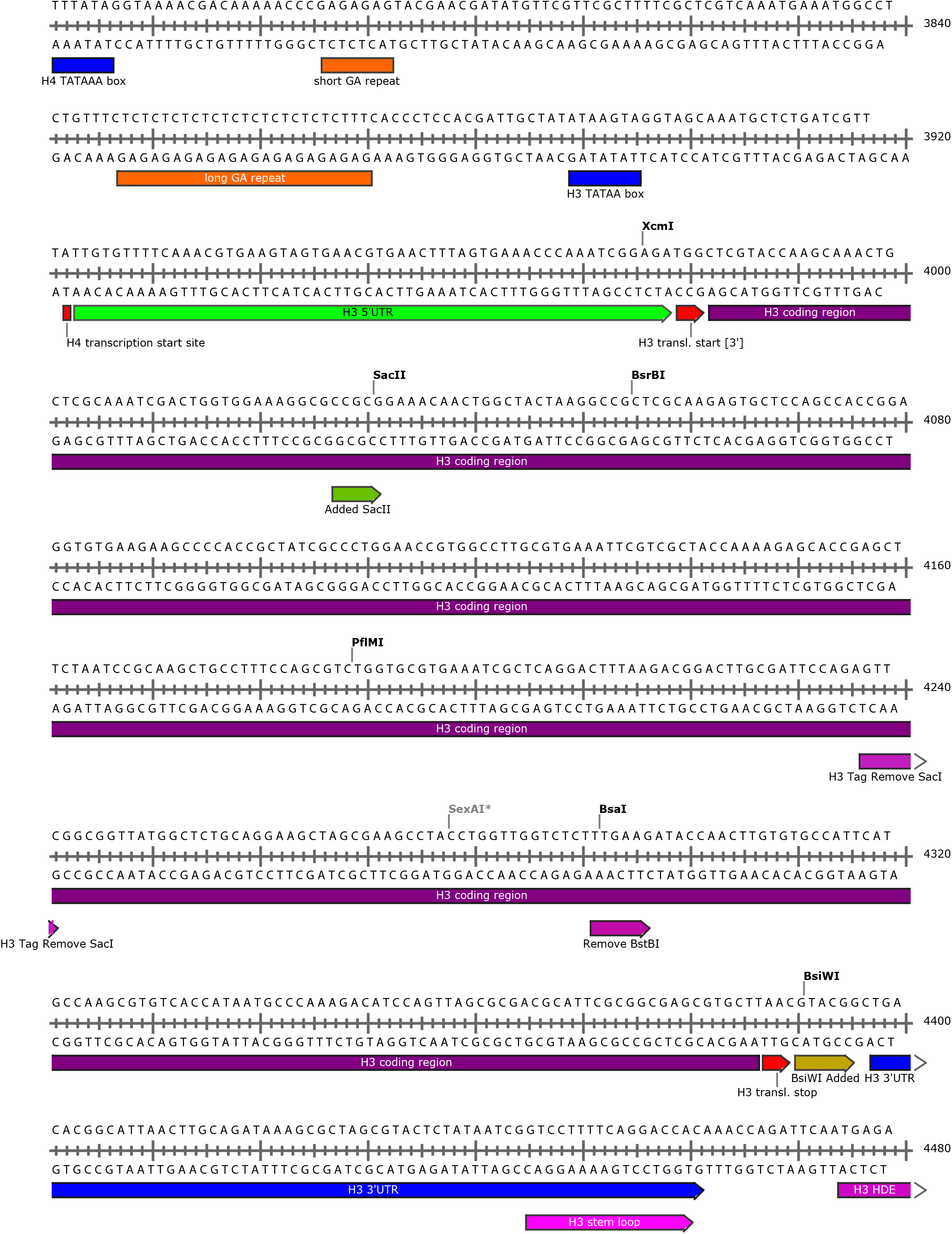
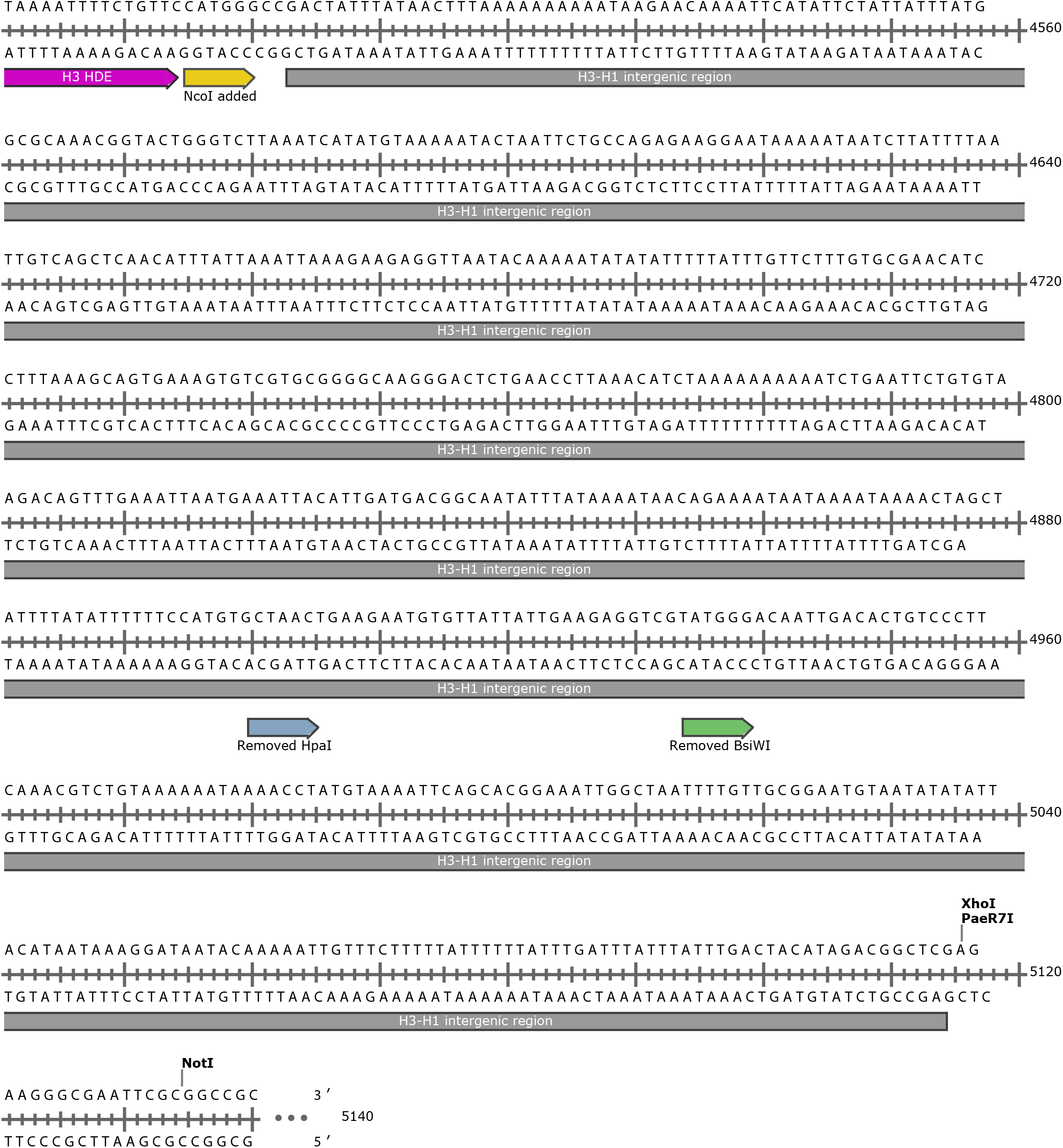
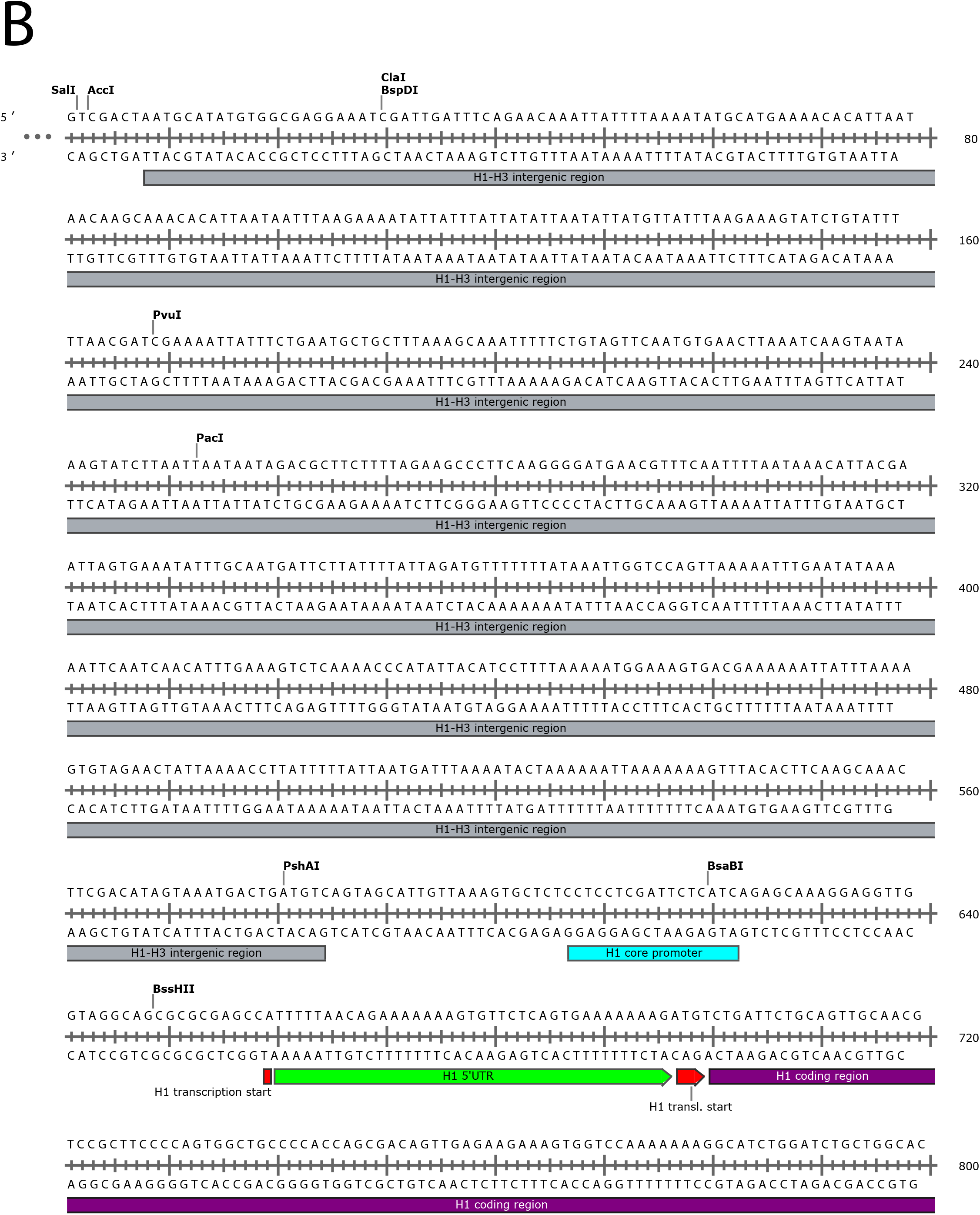
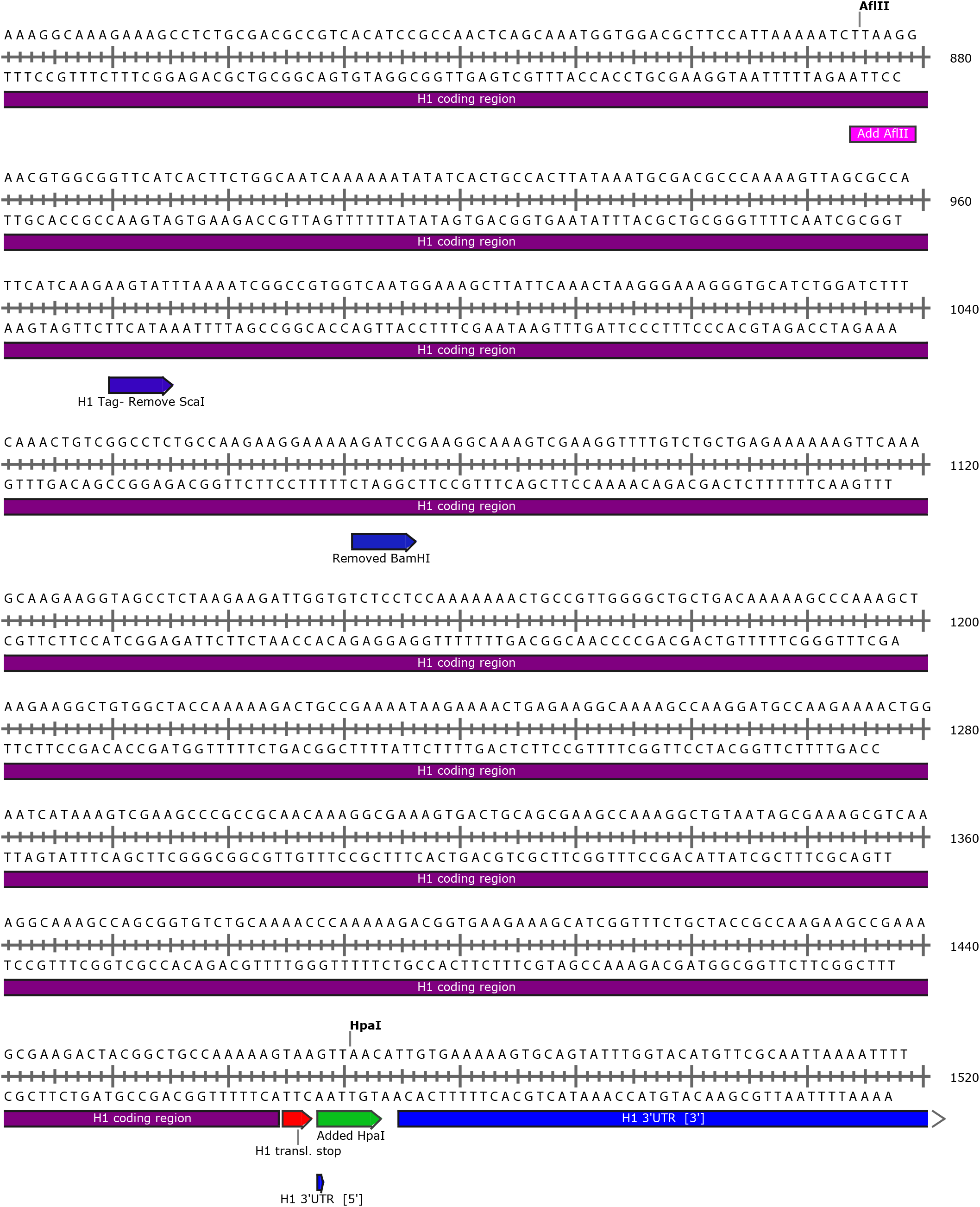
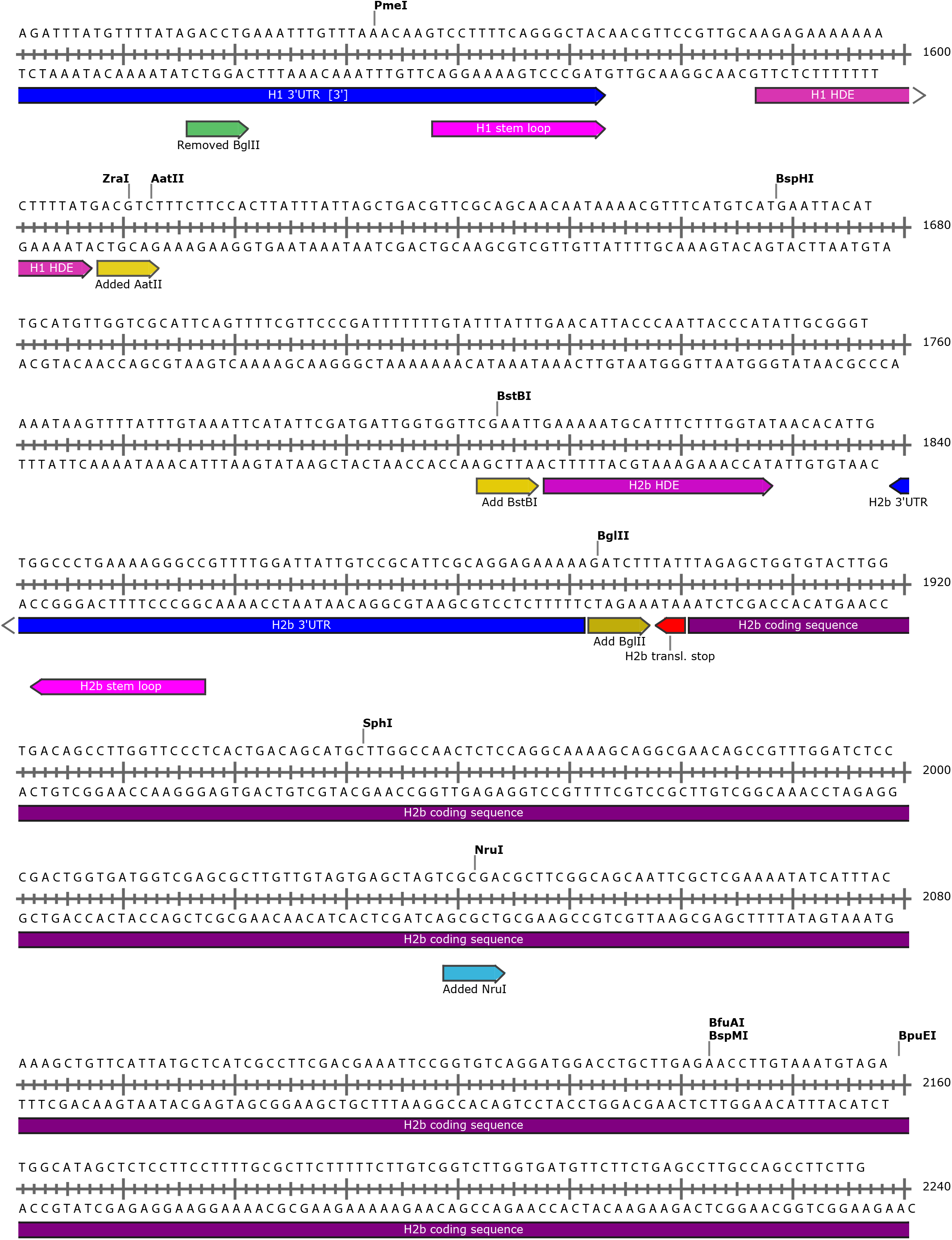
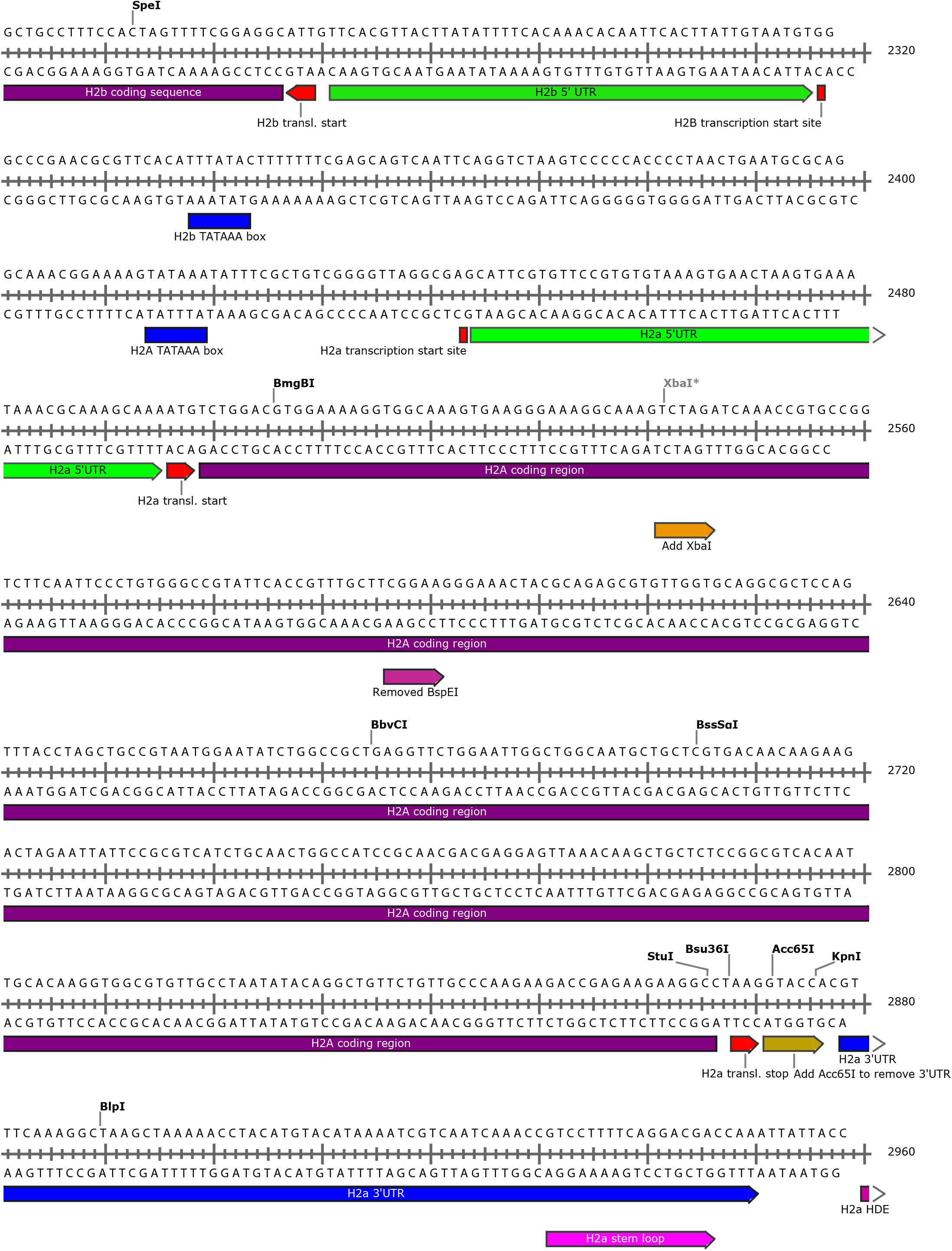
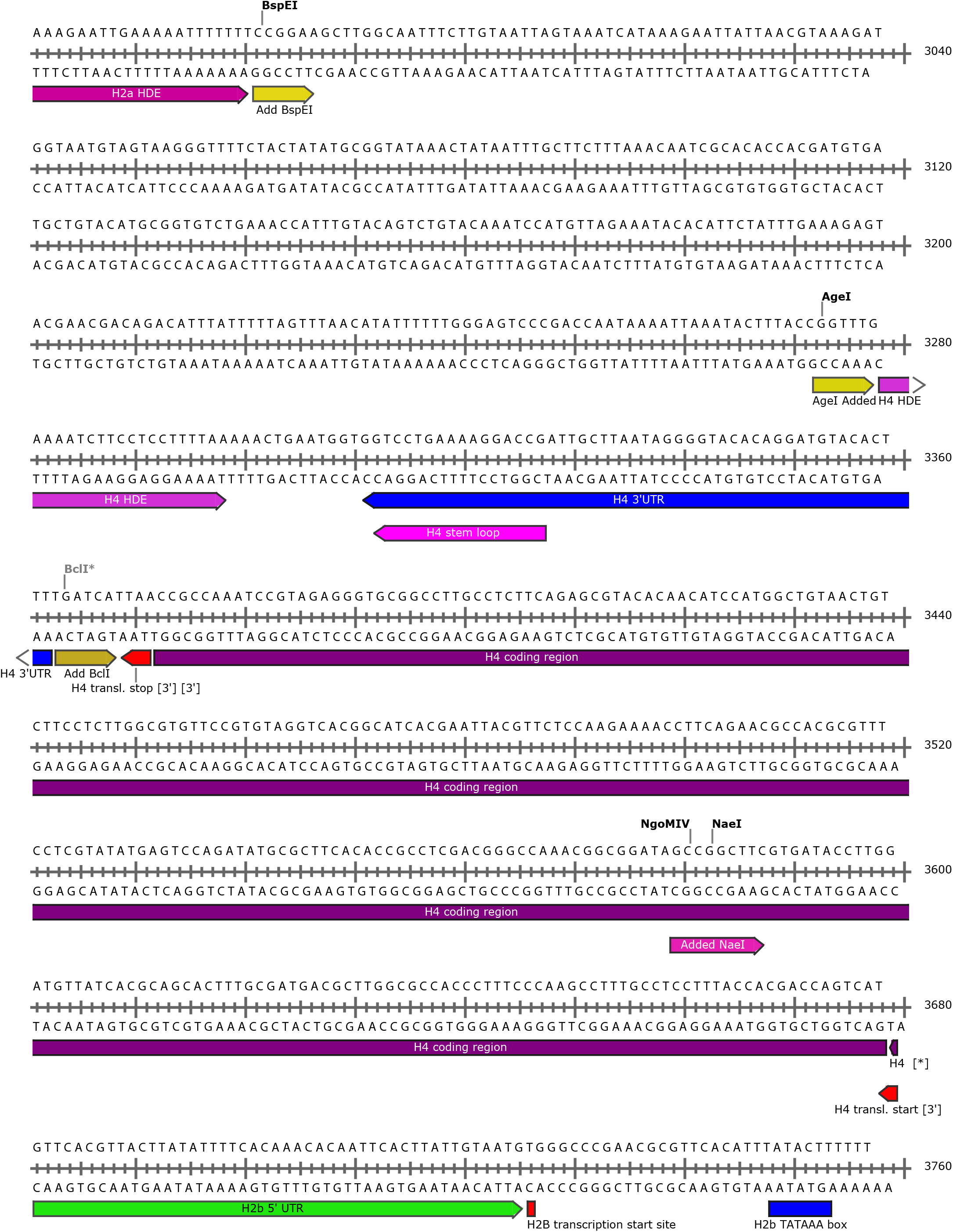
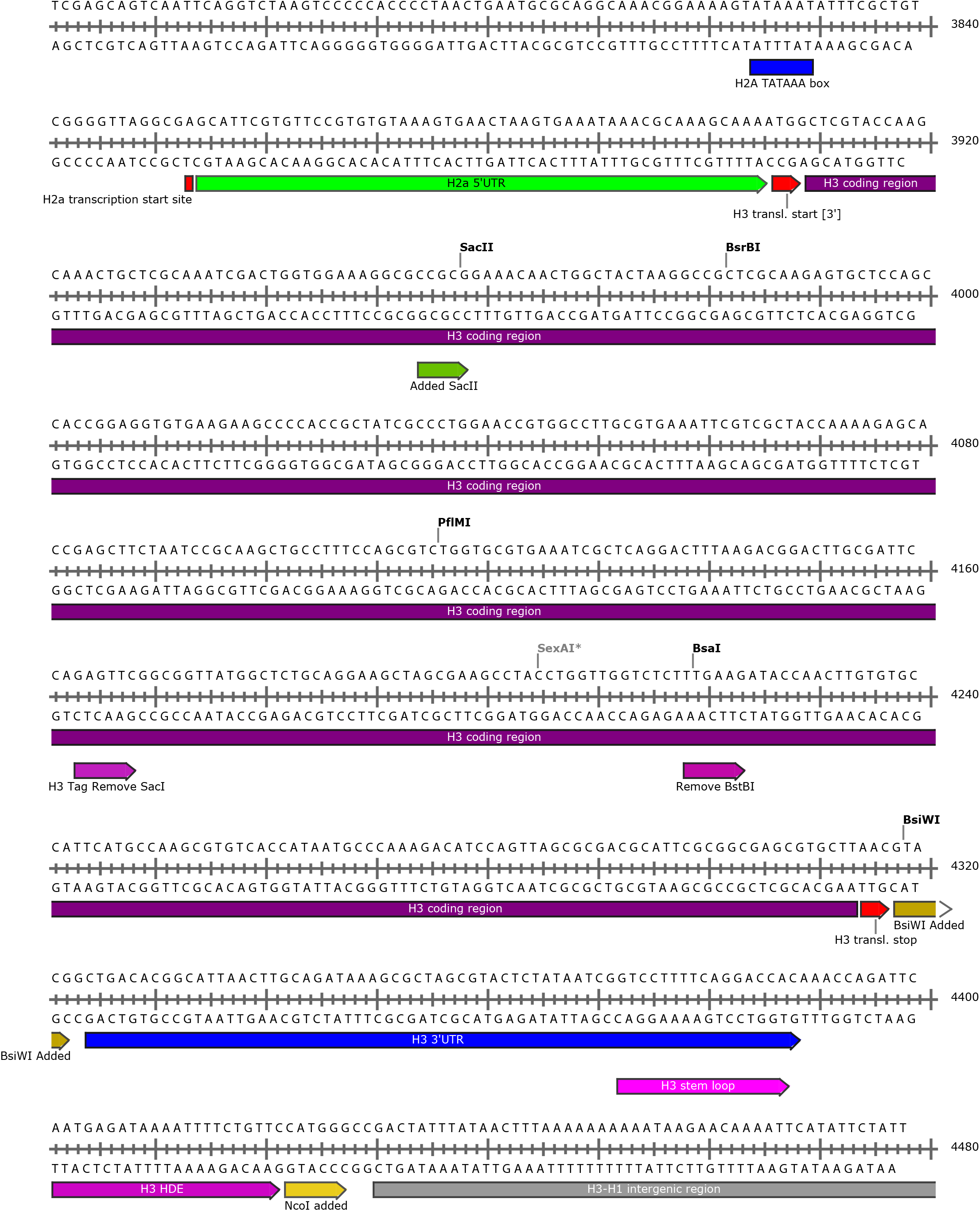
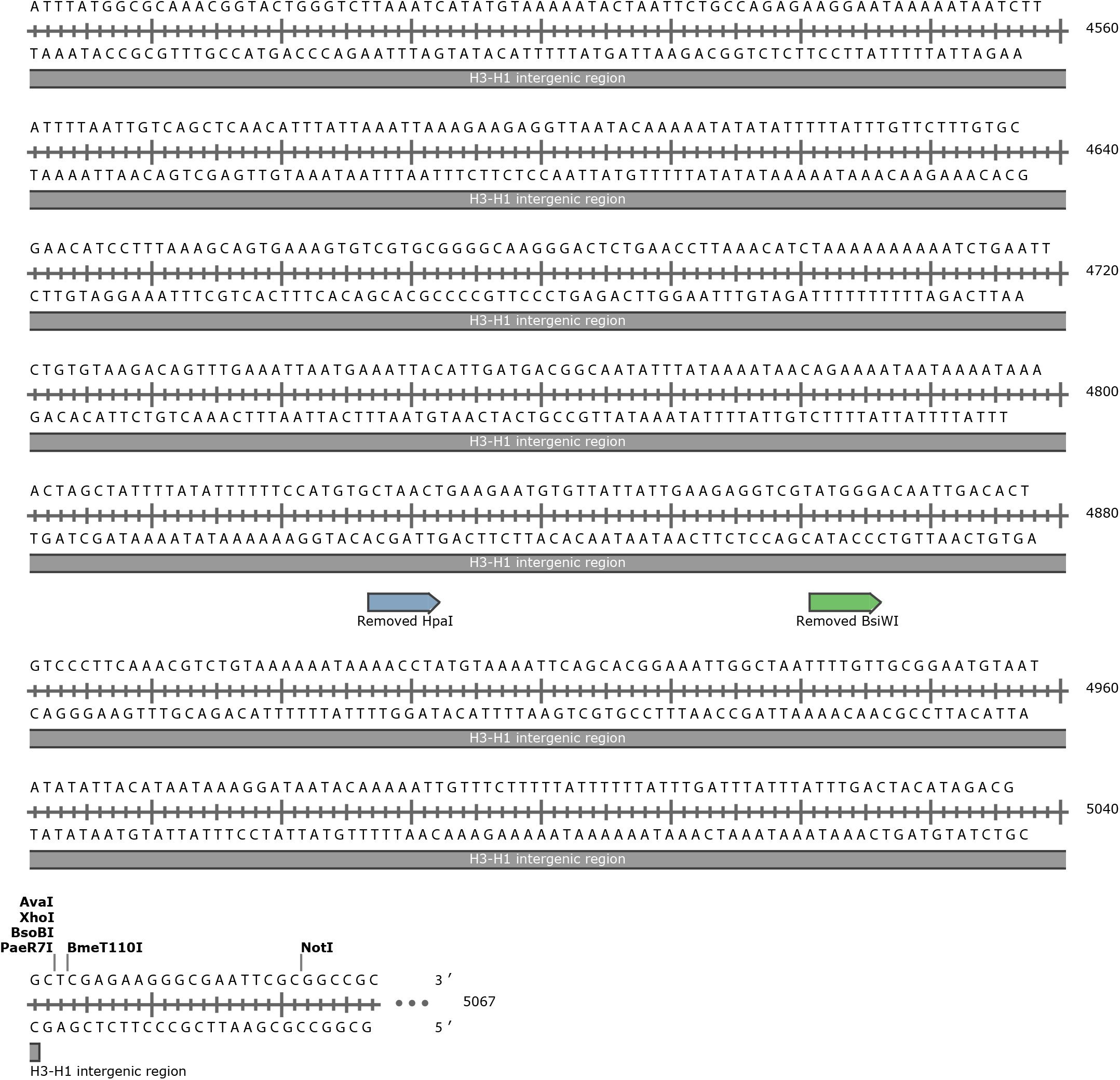
Annotated maps of engineered histone gene repeat unit used to make transgenic 12x replication dependent histone gene arrays. A) Designer Wild Type. B) Promoter Replacement. Restriction enzyme sites that were either added or deleted are shown below each line of sequence. The maps were made using Snapgene.

